# The coordination of terminal differentiation and cell cycle exit is mediated through the regulation of chromatin accessibility

**DOI:** 10.1101/488387

**Authors:** Yiqin Ma, Daniel J. McKay, Laura Buttitta

**Affiliations:** University of Michigan, Dept. of Molecular Cellular and Developmental Biology, Ann Arbor, MI; University of North Carolina, Chapel Hill, Dept. of Biology, Dept. of Genetics, Integrative Program for Biological and Genome Sciences, Chapel Hill, NC

## Abstract

During terminal differentiation most cells will exit the cell cycle and enter into a prolonged or permanent G0. Cell cycle exit is usually initiated through the repression of cell cycle gene expression by formation of a transcriptional repressor complex called DREAM. However when DREAM repressive function is compromised during terminal differentiation, additional unknown mechanisms act to stably repress cycling and ensure robust cell cycle exit. Here we provide evidence that developmentally programmed, temporal changes in chromatin accessibility at a subset of critical cell cycle genes acts to enforce cell cycle exit during terminal differentiation in the *Drosophila melanogaster* wing. We show that during terminal differentiation, chromatin closes at a set of pupal wing enhancers for the key rate-limiting cell cycle regulators *cycE*, *e2f1* and *stg*. This closing coincides with wing cells entering a robust postmitotic state that is strongly refractory to cell cycle re-activation. When cell cycle exit is genetically disrupted, chromatin accessibility at cell cycle genes remains largely unaffected and the closing of enhancers at *cycE*, *e2f1* and *stg* proceeds independent of the cell cycling status. Instead, disruption of cell cycle exit leads to changes in accessibility and expression of a subset of hormone-induced transcription factors involved in the progression of terminal differentiation. Our results uncover a mechanism that acts as a cell cycle-independent timer to limit aberrant cycling in terminally differentiating tissues. In addition, we provide a new molecular description of the cross-talk between cell cycle exit and terminal differentiation during metamorphosis.

## Introduction

The majority of cells in mature multicellular organisms spend most of their existence in non-proliferating states, often referred to as cellular quiescence or the G0 phase (O’Farrell, 2011). Substantial progress has been made on understanding how developmental signaling pathways interface with the cell cycle machinery to control tissue growth and proliferation (Ruijtenberg and van den Heuvel, 2016; Soufi and Dalton, 2016). Yet, we understand very little about why some cell types can enter a more flexible G0 state and retain the ability to re-enter the cell cycle, while others become permanently postmitotic. Robust and synchronous silencing of cell cycle gene expression is critical to the proper timing of cell cycle exit and the maintenance of a postmitotic state. Yet the molecular details of how this silencing is initiated and maintained in maturing tissues remains unresolved. This impacts a wide range of biological questions, as the proper control of G0 is critical during development and tissue regeneration, but becomes disrupted in cancer.

The transition from proliferation to G0 is accompanied by a functional switch in the master regulators of the cell cycle program, such as E2F transcription factor complexes, leading to global downregulation of cell cycle gene transcription (Blais and Dynlacht, 2007; Sadasivam and Decaprio, 2013; van den Heuvel and Dyson, 2008). In proliferating cells activating E2F family members (dE2F1 in *Drosophila*) binds with the Dimerization Partner, DP, at promoter proximal E2F binding motifs at hundreds of cell cycle genes, including cyclins, Cyclin-dependent kinases (Cdks), replication proteins and mitotic regulators, to promote their expression. Upon entry into G0, silencing through these same binding sites occurs via the formation of a transcriptional repressor complex called DREAM. DREAM complexes are conserved from *C. elegans* to humans and in *Drosophila*, DREAM (termed dREAM) consists of the E2F binding partner DP, RB family members Rbf1 or Rbf2, the repressive E2F transcription factor family member dE2F2, p55/CAF1, Myb and Myb-interacting proteins (Sadasivam and Decaprio, 2013; van den Heuvel and Dyson, 2008). Formation of dREAM is promoted by the accumulation of hypo-phosphorylated or unphosphorylated RB through the inhibition of cyclin/Cdk. Therefore it can be induced through developmental activation of Cyclin-dependent kinase inhibitors, developmental downregulation of the expression and production of cyclins and cdks or the upregulation of cyclin destruction through the APC/C (Buttitta et al., 2010; Buttitta et al., 2007; Firth and Baker, 2005; The et al., 2015).

While dREAM plays an important role in the repression of cell cycle genes in G0, key aspects of cell cycle exit *in vivo* are still not understood. For example, in some contexts of differentiation, cells eventually arrest and differentiate even in the presence of constitutive high E2F or Cyclin/Cdk activity (Camarda et al., 2004; Cecchini et al., 2014; Ebelt et al., 2008; Korzelius et al., 2011; Pajalunga et al., 1999). We and others have found that this is in part due to cooperative roles for RBs, Cyclin Kinase inhibitors and the APC/C. However, double and triple combinations of alleles altering these pathways cooperate to further delay cell cycle exit, but fail to abrogate it completely, suggesting these pathways act in addition to still unknown developmental mechanisms (Boxem and van den Heuvel, 2001; Buttitta et al., 2010; Di Stefano et al., 2011; Fay et al., 2002; Firth and Baker, 2005; van Rijnberk et al., 2017). These redundant mechanisms make cell cycle exit upon terminal differentiation more robust than other states of G0 such as reversible quiescence.

We and others have also observed that the longer a terminally differentiated cell remains in G0 the more refractory it becomes to re-entering the cell cycle, even in the presence of high E2F or Cyclin/Cdk activity (Coller et al., 2006). This has been termed “deep” or “robust” G0 (Buttitta et al., 2010; Yao, 2014). The molecular basis of robust G0 and how it differs from temporary or “flexible” G0 states remains unknown. One model for how terminally differentiated cells become resistant to strong proliferation signals involves a chromatin lockdown mechanism, where chromatin compaction or repressive modifications act globally to silence cell cycle gene expression and promote robust cell cycle exit. For example, DREAM complexes can recruit chromatin modifiers to add repressive histone modifications at E2F-dependent cell cycle genes such as H3K27 trimethylation (H3K27Me3) or H3K9 trimethylation (H3K9Me3) which in turn recruit repressive heterochromatin binding proteins such as the Polycomb complexes PRC1 or Heterochromatin Protein 1 (HP1) for long-term silencing of cell cycle genes (Blais et al., 2007; Sdek et al., 2011). Another model posits that cell cycle genes become recruited to the nuclear periphery to be sequestered in repressive nuclear lamina-associated domains (LADs) (Peric-Hupkes et al., 2010). We directly tested these models in the terminally differentiating *Drosophila* wing and found cell cycle exit occurs despite disruption of heterochromatin-dependent gene silencing (Ma and Buttitta, 2017) and without obvious sequestration or recruitment of cell cycle genes to heterochromatin compartments or the nuclear lamina. This suggests developmentally controlled cell cycle exit in *Drosophila* uses additional mechanisms to ensure a robust G0.

Cell cycle exit in the *Drosophila* wing occurs during metamorphosis and is tied to pulses of the hormone ecdysone that induce downstream transcription factors to modulate cell cycle gene expression (Guo et al., 2016). We further showed that transcription factors downstream of ecdysone signaling play a critical role in promoting sequential changes in chromatin accessibility to promote wing differentiation (Uyehara et al., 2017). This suggested to us that changes in chromatin accessibility during metamorphosis could contribute to the regulation of cell cycle genes to coordinate cell cycle exit with differentiation. To examine this, we characterized the transcriptome and genome-wide chromatin accessibility landscape of the *Drosophila* wing during metamorphosis through RNA-seq and FAIRE-seq over six developmental time points. We show that during wing differentiation, chromatin accessibility and gene expression changes are coordinated with the transition from a proliferating to a postmitotic state. This includes the closing of specific regulatory elements at a subset of critical “master” cell cycle regulators during G0. Moreover, we have uncoupled differentiation from cell cycle exit, revealing that the closing of enhancers at these cell cycle master regulator genes is developmentally programmed and occurs independent of E2F activity or cell cycling status, coincident with robust G0. We propose that the developmentally programmed closing of regulatory elements at a subset of key cell cycle genes is the molecular mechanism underlying robust cell cycle exit *in vivo*.

## Results

### Chromatin accessibility and gene expression are temporally dynamic during wing metamorphosis

During metamorphosis wings undergo morphogenetic changes coordinated with cell cycle alterations, loss of regeneration capacity and activation of a wing terminal differentiation program (Halme et al., 2010; Johnson and Milner, 1990; Schubiger et al., 2005). These events are temporally coordinated by systemic hormone pulses which trigger metamorphosis and drive its progression, leading to coordinated morphogenesis and differentiation of organs (Ashburner, 1990; Thummel, 2002). Although the hormone pulses are systemic, through a combination of direct and indirect regulation they result in activation of unique gene expression programs in different tissues (Ashburner, 1990; King-Jones and Thummel, 2005; Stoiber et al., 2016; Uyehara et al., 2017). For the wing, major events during metamorphosis include: eversion coordinated with a temporary cell cycle arrest in G2 and pupa cuticle formation, elongation and apposition of dorsal and ventral surfaces, coordinated with a relatively synchronized final cell cycle and vein refinement and finally permanent cell cycle arrest which precedes wing hair formation and deposition of adult cuticle (Fristrom and Liebrich, 1986; Guo et al., 2016; Sobala and Adler, 2016; Sotillos and De Celis, 2005; Taylor and Adler, 2008). Underlying these processes are temporally coordinated changes in gene expression. We and others have examined the dramatic gene expression changes in the wing during metamorphosis (O’Keefe et al., 2012; Sobala and Adler, 2016). To identify the global landscape of potential regulatory elements driving these gene expression changes, we carried out Formaldehyde-Assisted Isolation of Regulatory Elements sequencing (FAIRE-seq) in parallel with RNA-seq on a timecourse of wildtype *Drosophila* wings from the late wandering third instar stage when wing cells are proliferating to 44h after puparium formation (APF), when wing cells are postmitotic and begin to deposit adult cuticle.

We identified a total of 20,329 high-confidence open chromatin regions (peaks). We first compared the similarity of open chromatin profiles across our wing developmental time course by examining Pearson correlation coefficients. The open chromatin landscape is gradually changing during metamorphosis and early proliferative stages are clearly distinct from postmitotic stages in chromatin accessibility (Fig. 1A). By calculating the fold change in peak accessibility between stages, we found that only 5,516 peaks (27%) are static and exhibit <2-fold changes between any two timepoints. The remaining 14,813 peaks (73%) appear developmentally dynamic, exhibiting >2-fold changes between two or more timepoints. To visualize peak accessibility dynamics during metamorphosis, we divided the rpkm value of each FAIRE peak by its maximum rpkm value for each of the 6 timepoints, and then plotted the fraction in the form of heatmap (Fig. 1B). To distinguish different dynamic patterns, we separated the peaks into 18 k-means clusters. We found that dynamic peaks could be divided into 3 broad categories: a temporally sharp category that transiently opens at only one stage; a temporally broad category that remains accessible for several sequential stages and a category of peaks that oscillate during metamorphosis.

**Fig. 1.**
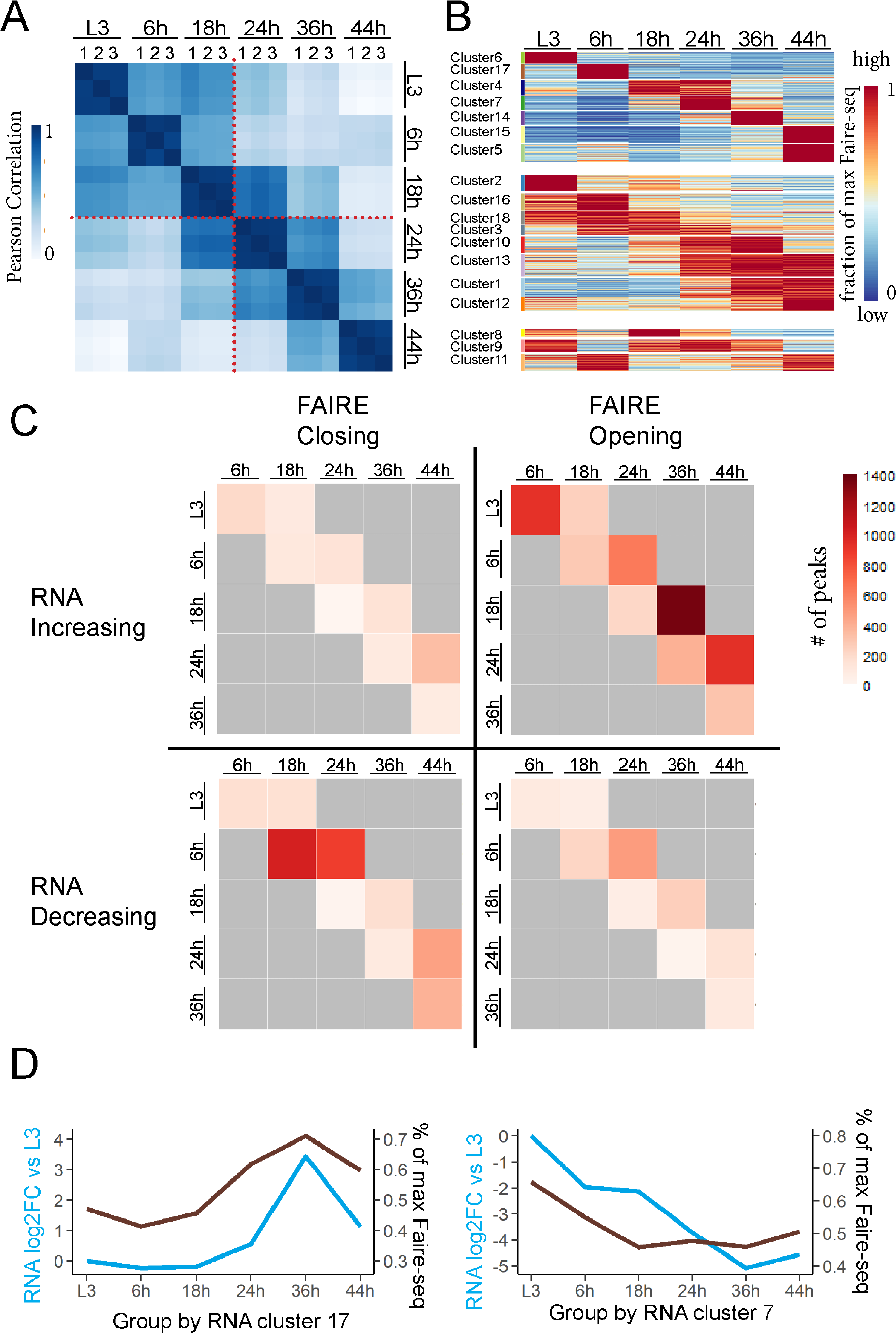
Dynamics of chromatin accessibility and gene expression are correlated during metamorphosis. (A) Open chromatin regions (peaks) in wings were identified by FAIRE-Seq on timepoints prior to metmorphosis (L3) and during pupal stages 6h, 18h, 24h, 36h and 44h APF. Heatmaps of Pearson correlation coefficients for each replicate across this timecourse reveal differences between the proliferative and postmitotic stages (red dotted line). (B) Dynamic open chromatin peaks were organized into 18 k-means clusters, displayed as a heatmap representing the fraction of the maximum FAIRE rpkm value. (C, D) Most chromatin accessibility changes are associated with gene activation rather than repression during metamorphosis. (C) We assigned dynamic FAIRE peaks to the nearest expressed gene and correlated peak changes (opening or closing) with observed gene expression changes (increasing or decreasing) measured by RNAseq at each subsequent timepoint. This revealed four classes of FAIRE peak/RNA expression correlations; opening/increasing consistent with gene activation, closing/decreasing consistent with loss of activation; opening/decreasing consistent with binding of a repressor and closing/activation consistent with a loss of repression. We show the number of dynamic FAIRE peaks that fall into each quadrant. (D) Genes were clustered based on RNA expression patterns across metamorphosis. Two clusters showing a high positive correlation between RNA signal (average log2 fold change from L3) and accessibility of their assigned FAIRE peaks (average maximum FAIRE rpkm value) are shown. The full dataset correlating RNA expression with accessibility of their assigned FAIRE peaks for all clusters is provided in the supplement.

Consistent with previous work, our parallel RNA-seq revealed dynamic expression changes for over 6,000 genes (over 35% of the genome) during wing metamorphosis. For comparison to the FAIRE-Seq clusters, we also clustered genes based on RNA expression into 18 k-means RNAseq clusters (Supp. Fig. 1A). Clustering based on gene expression identified groups of genes that are functionally related and temporally coordinated. For example, RNA cluster 4 contains genes highly expressed at 6h which are enriched for genes involved in wing pupal cuticle development (Supp. Fig. 1B) [32], while RNA clusters 7 and 10 coordinately decrease expression after 18h, and are highly enriched for cell cycle genes. This is consistent with our previous work showing that cell cycle gene expression plummets by 24h when cells transition to a postmitotic state (Guo et al., 2016; O’Keefe et al., 2012).

To more easily visualize the temporal dynamics of peaks, we next compared dynamic peaks between adjacent stages to define them as opening or closing compared to the previous stage (Fig. 1C-D, Supp. Fig. 2). The timepoint with the most dynamic changes is 6h while the second most dynamic is 24h. Both of these stages are associated with cell cycle arrests. We previously showed that at 6h wings undergo a temporary G2 arrest induced by high levels of the transcription factor Broad suppressing the critical G2-M regulator cdc25c or string. This synchronizes the subsequent final cell cycle (Guo et al., 2016). At 24h cells in the wing finish the final cell cycle in a relatively synchronized manner and enter into a postmitotic G0 state (Milan et al., 1996; Schubiger and Palka, 1987). This suggested to us that a developmentally controlled program of coordinated chromatin accessibility could link changes in the cell cycle with differentiation during metamorphosis.

**Fig. 2.**
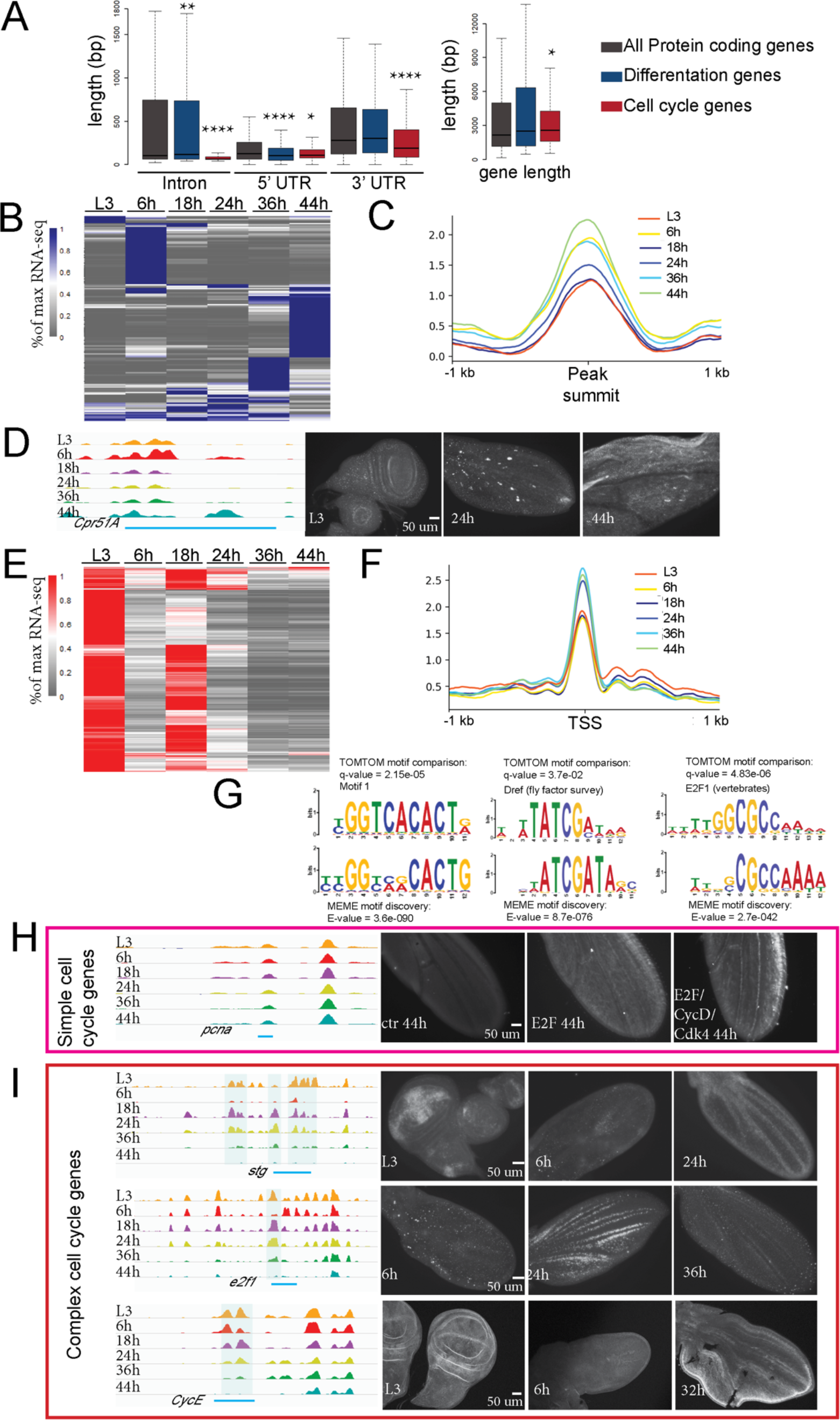
Temporal regulation of the wing differentiation program and cell cycle changes. (A) The length (in bp) of introns, 5’, 3’ UTRs (right) and genes (left) for all protein coding genes, wing terminal differentiation genes and cell cycle genes is shown. The majority of FAIRE peaks occur within introns (Supp Fig. 2). Most cell cycle genes have a compact structure with few, short introns, while differentiation genes contain large introns, providing potential dynamic regulatory elements. (B, E) Heatmap of gene expression for differentiation genes (B) and cell cycle genes (E), plotted by the % of maximum RNA rpkm value. Both groups of genes show dynamic expression during metamorphosis. (C, F) Line plots of average FAIRE signal of the 6 stages for differentiation genes (C) and cell cycle genes (F). Differentiation genes show an increase in FAIRE peak accessibility at timepoints when gene expression is high: 6h (p-value = 0.0004088), 36h (p-value = 1.36e-07) and 44h (p-value = 1.408e-12), compared to L3, Mann-Whitney U Test). Cell cycle genes show an increase in accessibility at timepoints when gene expression is repressed: 24h (p-value = 0.0209), 36h (p-value = 1.655e-05), 44h (p-value = 0.005469), Mann-Whitney U Test. (D) A Gal4 reporter containing the indicated (blue line) portion of the Cpr51A regulatory region drives UAS-GFP in late wings (44h) when the regulatory elements are accessible. (G) Motif discovery was performed on FAIRE peaks for cell cycle genes using MEME and compared to known motifs using TOMTOM. Potential regulatory elements for cell cycle genes are highly enriched for E2F binding motifs, DRE promoter sequences and the Pol II pausing-associated motif1. (H) A GFP reporter containing the indicated regulatory element for the simple cell cycle gene, *pcna* is silent at the postmitotic stage of 44h, but can be re-activated postmitotically when E2F or E2F+CycD/Cdk4 is expressed. (I) *Stg*, *e2f1* and *cycE* are complex cell cycle genes with large dynamic regulatory regions. Gal4 reporters containing the indicated portions of their regulatory regions drives UAS-degradable-GFP to capture their regulatory dynamics. Expression correlates with accessibility for these regions. P-values were determined by Mann-Whitney U Test; **** <0.0001, **<0.01, *<0.05

To correlate chromatin accessibility changes with gene expression changes, we assigned dynamic and static FAIRE peaks to the nearest transcription starting site (TSS) (Supp. Fig. 2). Dynamic and static FAIRE peaks exhibit similar distributions, with most of them located in introns, intergenic and promoter proximal regions. This is consistent with previous work showing that FAIRE-seq enriches for DNA regulatory elements (Song et al., 2011). Both dynamic and static peaks were most highly enriched at locations near (1-5 kb) their assigned TSS. However dynamic peaks were also more likely to be located further from TSSs (5-10 kb) than static peaks, which were more likely to be promoter proximal (within 0.5 kb from the assigned TSS) (Supp. Fig. 2). Similar to other studies using FAIRE in *Drosophila*, we find most developmentally dynamic putative regulatory elements for the wing are located within introns, especially the first intron and1-5 kb upstream of the TSS. This is also consistent with the locations of *Drosophila* enhancers identified using a functional accessibility-independent approach, Starr-Seq (Arnold et al., 2013).

### Dynamic chromatin is mostly correlated with gene activation

Open chromatin sites often correspond to gene regulatory elements such as transcriptional enhancers which can activate or repress gene expression. To determine whether FAIRE peak dynamics correlate positively or negatively with gene expression changes, we assigned each FAIRE peak to its nearest gene and carried out pair-wise comparisons between each stage and its next two sequential stages for >2-fold changes in chromatin accessibility correlated with >2-fold expression changes using our RNAseq data (Supp. Fig. 3). When we plot peak accessibility change vs. assigned gene expression changes, we generate four quadrants: FAIRE peaks opening with corresponding gene expression increasing consistent with an activation function; FAIRE peaks opening with gene expression decreasing consistent with a repressive function; FAIRE peaks closing with gene expression increasing consistent with the loss of a repressor binding; FAIRE peaks closing and gene expression decreasing consistent with the loss of an activator binding (Fig. 1C).

We observed that the majority of dynamic FAIRE peaks fall into the category of peaks opening with the corresponding gene expression increasing. This suggests that the majority of gene expression changes in the differentiating wing are driven by transcriptional activators gaining access to their binding sites. The second largest category of FAIRE peaks close and are associated with loss of expression. This suggests that loss of access to transcriptional activators also plays a major role in gene repression during terminal differentiation.

We next examined the correlation between gene expression and chromatin accessibility changes during our timecourse (Fig. 1D, Supp. Fig. 4-5). We plotted the trajectory of gene expression based upon RNAseq for 18 co-regulated gene clusters and overlaid the average changes in FAIRE peaks assigned to the genes within each RNA cluster (Fig. 1D, Supp. Fig. 4).

We also performed a reciprocal analysis using 18 gene clusters based upon co-regulated FAIRE peaks and overlaid average gene expression changes from RNAseq (Supp. Fig. 5). We found that for several clusters the temporal changes in RNA and accessibility by FAIRE-seq are well correlated (Fig. 1C). Together our results suggest that most of the dynamic regulatory elements during fly wing metamorphosis are associated with gene activation.

### Opening the wing differentiation program during metamorphosis

A major event during wing differentiation is the formation of the adult cuticular exoskeleton. The wing cuticle is a multilayered structure and its formation requires the proper expression of cuticle-related proteins, such as enzymes involved in cuticle deposition, and ZP domain proteins which link the apical surface of wing cells to the overlying cuticle (Sobala and Adler, 2016). When we examined 154 cuticle formation-associated genes, we found distinct subgroups of genes highly expressed at 6h, 36 and 44h (Fig. 2B). The cuticle genes expressed at 6h are likely to be involved in the pupa cuticle formation (Fristrom and Liebrich, 1986), while the adult cuticle program begins at 36h and extends to 44h and beyond. The subgroups of cuticle-related genes reaching their peak expression at different stages suggests that waves of sequential regulation during metamorphosis may drive differences in pupa cuticle vs. adult cuticle composition and structure. Highly accessible FAIRE peaks found near cuticle genes (Fig. 2C) are significantly more accessible at 6, 36 and 44h, consistent with the high expression at those time points. To identify a cuticle gene enhancer, we examined a line containing a Gal4 transgene overlapping an open chromatin region near the cuticle gene *Cpr51A* driving UAS-GFP (Fig. 2D). This region is highly accessible at 44h and with this transgene GFP is highly expressed in almost all the wing cells at 44h. Opening and activation of the adult cuticle program is a major feature of wing differentiation during metamorphosis.

### Repression of most cell cycle genes is established and maintained through promoter proximal regulatory elements

Cells in pupal wings exit the cell cycle at 24h, which accompanies temporally synchronized repression of cell cycle genes. We examined the expression of ~300 cell cycle genes compiled from our previous analysis of cell cycle exit (Buttitta et al., 2010) (Fig. 2E) and observed a temporary repression during the G2 arrest at 6h, followed by reactivation at 18h for the final cell cycle, and silencing during cell cycle exit at 24h. We examined the chromatin accessibility profiles for 291 of these cell cycle genes and found that most of them exhibit a compact gene structure with smaller introns and relatively short intergenic upstream sequence (Fig. 2A). Most FAIRE peaks associated with these genes are found to be proximal to the TSS, consistent with the previously reported distribution for functional enhancers at “housekeeping” genes (Zabidi et al., 2015). Surprisingly, putative regulatory elements at cell cycle genes exhibit a moderate increase in accessibility at timepoints when cells are postmitotic (24-44h) despite the temporally regulated shutoff of their associated genes at 24h (Fig. 2F).

We carried out a *de novo* motif discovery on the promoter proximal FAIRE peaks for cell cycle genes using MEME (Fig. 2G). The most highly enriched motifs match Motif 1, a core promoter element bound by M1BP to promote RNAPolII pausing (Li and Gilmour, 2013; Ohler et al., 2002), the Dref-binding element DRE, a core promoter/enhancer known to be associated with cell cycle genes (Matsukage et al., 2008) and a motif matching the binding site for the heterodimer transcription factor complex E2F/DP. The increased accessibility at these motifs is similar to the increased MNase sensitivity found at sites in cell cycle genes bound by repressive human DREAM complexes (Marceau et al., 2016). While increased accessibility may seem counter-intuitive when coupled with gene repression, this has been suggested to be consistent with a model whereby promoter proximal DREAM binding to nucleosome free regions represses cell cycle genes by positioning nucleosomes downstream of the transcriptional start site (Marceau et al., 2016).

Multiple studies have reported that depletion of Rb family members leads to de-repression of cell cycle genes and defects in exiting the cell cycle. However it has remained unclear whether Rb- or DREAM-dependent repression is required to counteract E2F activity to initiate repression of cell cycle genes, or to maintain repression in cells that have already become postmitotic, or both. To investigate this we took advantage of a PCNA-GFP transcriptional reporter that includes known E2F binding sites contained within FAIRE peaks that remain accessible after cell cycle exit (Thacker et al., 2003). At 44h, a timepoint when the postmitotic state has been maintained for 20h, the reporter is silenced. To test whether this silencing can be reversed, we activated expression of the *Drosophila* activator E2F complex E2F1/DP (hereafter E2F) or E2F+CycD/Cdk4 to phosphorylate and inactivate Rbf specifically after cells have already established a flexible G0 state at 26h (Fig. 2H). Expressing either E2F or E2F/CycD/Cdk4 was able to re-induce PCNA-GFP expression in postmitotic cells, demonstrating that RB/E2F-dependent repression is required to maintain silencing of cell cycle genes in *Drosophila*.

### The accessibility of enhancers for complex cell cycle genes are dynamic

In contrast to the majority of cell cycle genes, a few key, rate-limiting cell cycle genes are controlled by long, complex regulatory elements upstream of their TSS or in long introns. For example, *cycE*, *stg* and *e2f1* fall into this group (Fig. 2I). We find several FAIRE peaks in regulatory regions for these genes that overlap with previously characterized functional regulatory elements (Andrade-Zapata and Baonza, 2014; Bradley-Gill et al., 2016; Jones et al., 2000; Lehman et al., 1999). Here we discovered that the accessibility of these regulatory elements is temporally dynamic during metamorphosis, in a manner coordinated with the cell cycle changes. Accessibility at these elements is low during the G2 arrest at 6h, then rises at 18h and 24h and closes after 36h. To examine whether the dynamic accessibility of these elements impacts temporal gene expression, we tested regions from the *stg*, *e2f1* and *cycE* loci driving a Gal4/UAS-destabilized GFP (*stg, e2f1*) or normal GFP (*cycE*) to capture gene expression shutoff. Our GFP reporters showed dynamic expression correlated with the accessibility of the elements and verifies these elements as pupal wing enhancers for these cell cycle genes. Our results suggest that dynamic chromatin accessibility at specific enhancers of complex cell cycle genes drives temporal expression changes during metamorphosis.

### The closing of enhancers at complex cell cycle genes is independent of cell cycle exit

We observed that chromatin dynamics at master regulator cell cycle genes is coordinated with cell cycle changes during metamorphosis. However, a key question is whether the closing of chromatin at these genes is a cause or consequence of cell cycle exit. To address this question, we took advantage of conditions where cell cycle exit in the pupal wing can be either temporarily delayed or bypassed for a prolonged period without preventing metamorphosis or terminal differentiation. In brief, overexpression of the activator E2F complex during the final cell cycle delays cell cycle exit and causes an extra cell cycle during the 24-36h window, while overexpression of E2F + CycD/Cdk4 during this same period causes multiple rounds of extra cell division and effectively bypasses cell cycle exit until well after 50h (Ma and Buttitta, 2017). We used the Gal4/UAS system in combination with a temperature-sensitive tub-Gal80 (Gal80^TS^) to limit the perturbation of the cell cycle from 12h −24 or 44h APF. This allows metamorphosis to initiate properly, yet effectively delays G0 by one extra cell cycle or bypasses G0 with multiple rounds of extra division (Fig. 3A, Supp. Fig. 6). We dissected 24h or 44h pupal wings under the delayed (E2F) or continued cycling (E2F+CycD) conditions and performed genome wide RNA-seq and FAIRE-seq analysis (Fig. 3B-C). Importantly, at 44h when the E2F expressing wings are postmitotic, the E2F+CycD wings are still cycling, allowing us to distinguish the effects of E2F overexpression from those of preventing cell cycle exit.

**Fig. 3.**
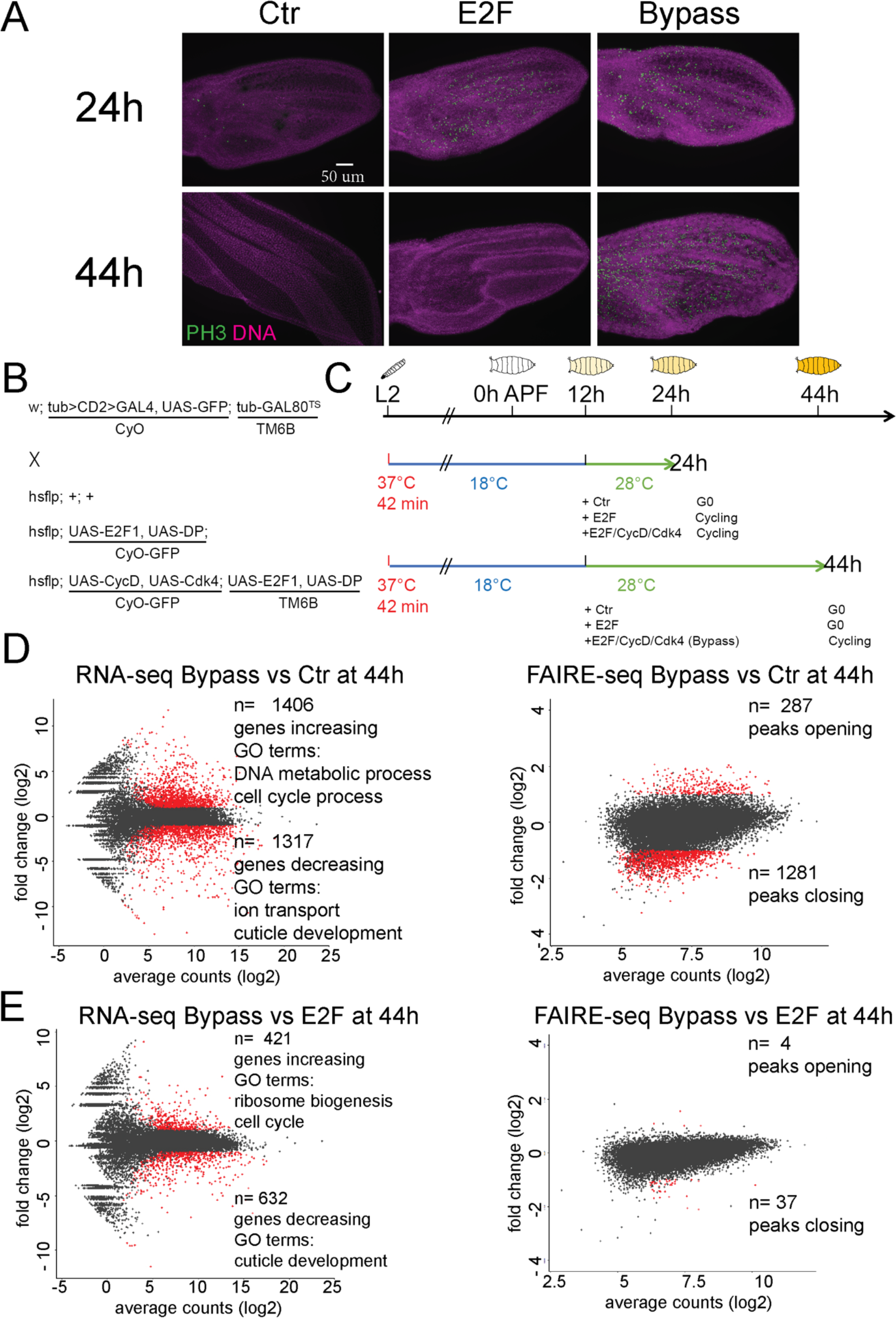
Global impacts of cell cycle exit disruption on gene expression and open chromatin. (A) G0 can be delayed to 36h or bypassed beyond 50h through short term expression of E2F or E2F/CycD/Cdk4. Transgenes were overexpressed in the dorsal layer of wing epithelia under the control of Apterous-Gal4/Gal80ts from 12h APF. 24h and 44h wings were immunostained against phospho-histone H3 (ph3). (B, C) Genotype and scheme of RNA-seq and FAIRE-seq experiments to disrupt cell cycle exit during metamorphosis. (D, E) MA-plots of RNA and FAIRE changes comparing bypassed exit (E2F+CycD/cdk4) between control (D) and delayed cell cycle exit (E2F) (E) at 44h. Abundant changes in expression of cell cycle genes, ribosome biogenesis and cuticle formation genes are observed, while chromatin accessibility is nearly identical between conditions where cells enter a delayed G0 vs continue cycling.

E2F or E2F+CycD expression was sufficient to alter the expression of several hundred genes at 24h and over 1,500 by 44h (Fig. 3D, Supp. Fig. 7A). Despite the dramatic changes in gene expression, there were strikingly few changes in FAIRE peak accessibility, with only a handful of peaks increasing accessibility at the 24h timepoint and up to 287 peaks increasing accessibility at the 44h timepoint (Fig. 3D, Supp. Fig. 7B). GO term enrichment analysis under both conditions revealed that the upregulated genes are highly enriched for those associated with the cell cycle while downregulated genes are highly enriched for genes involved in cuticle development. To determine whether these few accessibility changes were due to the ectopic E2F activity or the continued ectopic proliferation itself, we compared chromatin accessibility changes between E2F and E2F+CycD wings at 44h (Fig. 3F). While RNA-seq revealed differential effects on the expression levels of several hundred genes involved in the cell cycle, ribosome biogenesis and cuticle development, FAIRE-Seq revealed almost no changes in chromatin accessibility between these two conditions. This is remarkable considering that wings expressing E2F at 44h are fully postmitotic while wings expressing E2F+CycD continue to proliferate (Supp. Fig. 6). This demonstrates that the cycling status of differentiating cells has little direct effect on chromatin accessibility at potential regulatory elements.

Despite the upregulation of hundreds of cell cycle-related genes at both 24 and 44h (Fig.3 D,E, 4A), we observed little effect on their accessibility (Fig. 4B). Examples of simple (*orc6* and *pcna*) and complex cell cycle genes (*cyce* and *stg)* showed minor changes in chromatin accessibility when cell cycle exit was delayed or disrupted. Importantly, the closing of *cyce* and *stg* enhancers proceeds with normal timing despite the delay or bypass of cell cycle exit (Fig. 4C). This demonstrates that we have uncoupled differentiation from cell cycle exit and that the closing of enhancers at complex cell cycle genes is developmentally controlled and independent of cell cycling status. Importantly, the closing of enhancers prevents the activation of *stg* by ectopic E2F but not E2F + CycD (Fig. 4D). This suggests that the continued closing of enhancers at these cell cycle genes underlies the increased robustness of the G0 state at these later timepoints.

**Fig. 4.**
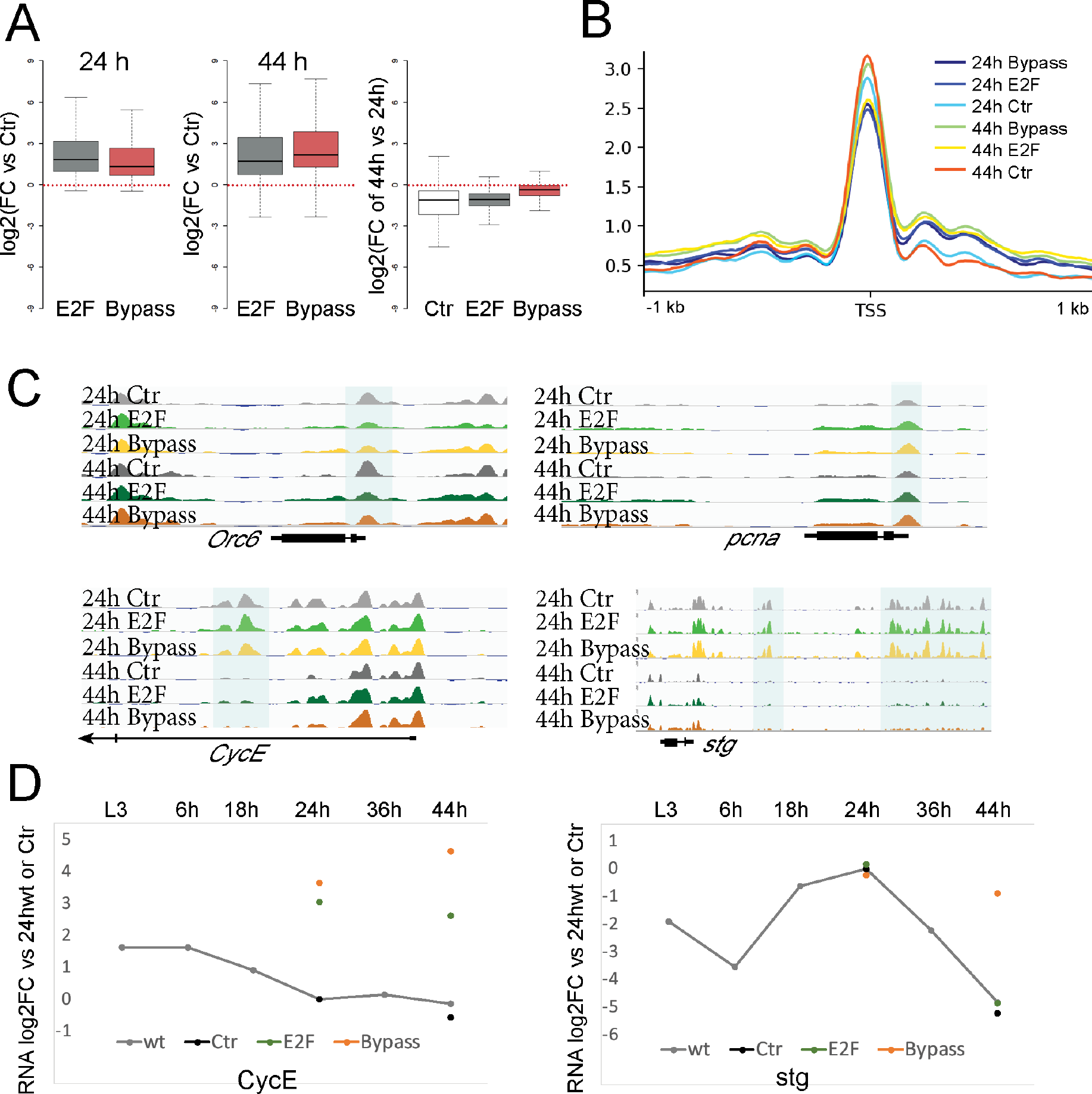
Enhancer accessibility of complex cell cycle genes is developmentally controlled and independent of cell cycling status. (A) Expression of cell cycle genes is increased when we delay or bypass cell cycle exit (log2 fold change for cell cycle genes vs. controls expressing GFP). (B) Line plots of average FAIRE signal for cell cycle genes. Accessibility at most cell cycle genes TSS is slightly decreased when cell cycle exit is delayed (44h E2F expression, p-value = 1.004e-05, Mann-Whitney U Test). (C) Regulatory elements for simple cell cycle genes (*orc6*, *pcna*) remain accessible independent of cycling status. Complex cell cycle genes (*cyce*, *stg*) lose accessibility at regulatory regions independent of cycling status. (D) Expression of *cycE* and *stg* during metamorphosis (gray line, compared to 24h wings) and genetic manipulations (colored dots, compared to 24h ctr wings). *stg* possesses higher barrier for activation compared to *CycE*. Closed stg regulatory elements prevent *stg* expression in the late robust E2F expressing wings, while E2F + CycD/Cdk4 can overcome this proliferation barrier.

### Delaying cell cycle exit impacts a subset of genes involved in wing terminal differentiation

In contrast to the minimal effects on cell cycle genes, the largest impact of delaying or disrupting cell cycle exit on chromatin was the loss of accessibility at over 1,000 genomic sites at 44h (Supp. Fig 7B). This could be caused by either chromatin remodeling to close accessible sites at 44h or a failure to open sites that should become accessible. To address which of these scenarios is correct, we examined the dynamics of peaks influenced by E2F or E2F+CycD during the wildtype time-course (Supp. Fig. 9A). Notably, peaks that are less accessible in E2F expressing wings are closed at 36h but highly accessible at 44h in wild type wings. This suggests that delaying or disrupting cell cycle exit results in a failure to open a specific subset of regulatory elements between 36h and 44h. Our data suggests that this failure to open specific elements is due to the ectopic E2F activity rather than ectopic proliferation itself, as there are strikingly few chromatin accessibility changes between E2F and E2F+CycD wings at 44h (Fig. 3E).

The loci that fail to open when cell cycle exit is disrupted are located near genes enriched for roles in cuticle formation and deposition and wing terminal differentiation. Consistent with this, expression levels of genes involved in wing cuticle formation are reduced when cell cycle exit is delayed or disrupted (Fig. 5A), and chromatin accessibility at their potential enhancers is reduced (Fig. 5B, C). Together, our results indicate that delayed cell cycle exit and ectopic E2F activity compromises the opening and activation of a portion of the wing terminal differentiation program.

**Fig. 5.**
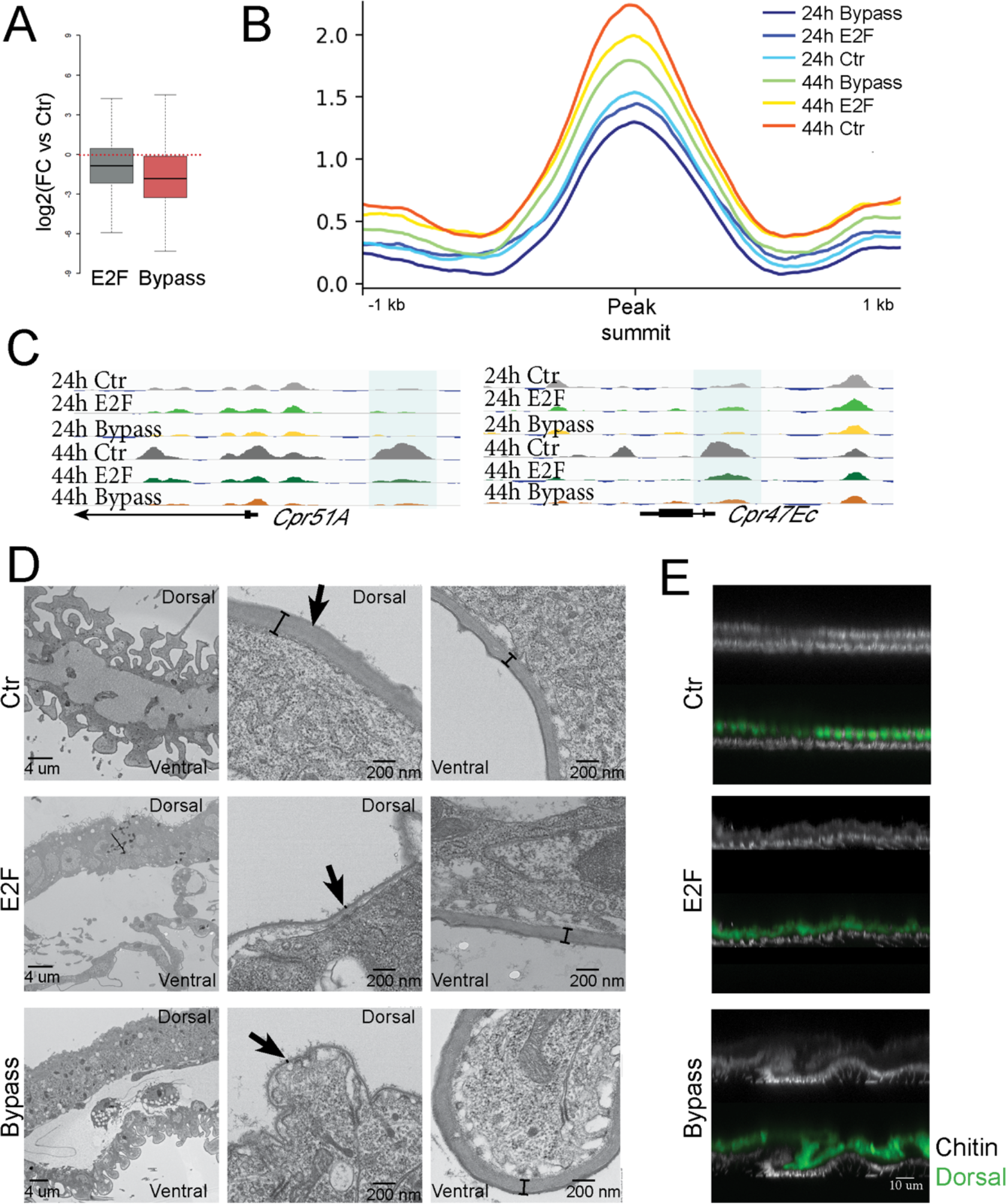
Compromising cell cycle exit impacts chromatin accessibility and gene expression at a subset of wing terminal differentiation genes. (A) log2 fold changes in RNA and (B) line plots of average FAIRE signal for genes involved in cuticle formation and differentiation. Preventing cell cycle exit reduces their expression and chromatin accessibility (E2F +CycD/Cdk4 at 44h, p-value = 0.004644, Mann-Whitney U Test). (C) Selected cuticle protein genes exhibiting a failure to open potential regulatory elements at 44h when cell cycle exit is delayed or bypassed. (D, E) TEM (D) and chitin staining (E) of 64h wings that delayed or bypass cell cycle exit in the dorsal wing epithelium using *Apterous*-Gal4/Gal80^ts^ to activate E2F or E2F+CycD/Cdk4 expression during the final cell cycle. Extra cellular matrix formation and chitin deposition are disrupted when cell cycle exit is compromised.

To determine whether ectopic E2F activity impacts wing cuticle formation, we expressed E2F or E2F+CycD in the dorsal layer of the wing epithelium beginning at 12h APF using *Apterous*-Gal4/Gal80^ts^. We examined the cuticle formation at 64h by transmission electron microscopy (TEM) (Fig. 5D). Pupal wings are normally composed of two thin monolayers of epithelial cells, and expression of E2F or E2F+CycD led to an obvious thickening of the epithelium due to extra cell divisions in the dorsal side. The cuticle layer on the dorsal side of the wing was much thinner than normal, and the effect on the cuticle was compartment autonomous, leaving ventral wing cuticle unaffected. We next examined the deposition of chitin, the key component of insect cuticle through calcofluor staining (Fig. 5E). Chitin staining in the dorsal wing where E2F or E2F+CycD was expressed was much weaker than the ventral. Thus, ectopic E2F activity delays and disrupts the adult wing cuticle program in a compartment autonomous manner.

### Disrupting cell cycle exit alters chromatin dynamics at specific ecdysone target genes

Our findings suggest the existence of crosstalk between cell cycle exit and the later terminal differentiation gene expression programs. We next sought to identify the factors mediating this crosstalk. We noticed that several ecdysone target genes were amongst the genes impacted by the delay or disruption of cell cycle exit (Supp Fig. 9A,C). Ecdysone signaling coordinates developmental timing between tissues during metamorphosis. We systematically examined ecdysone target genes and found that genes such as *Blimp-1*, *Hr3* and *crol* were expressed at significantly higher levels at 44h when cell cycle exit was disrupted while the expression of *E74EF*, *E75B* and *E71CD* was reduced (Fig. 6A). During the normal timecourse *Blimp-1*, *Hr3* and *crol* exhibit peak expression at 36h and plummet by 44h, while *E74EF*, *E75B* and *E71CD* normally peak at 44h. Thus the disruption of cell cycle exit leads to a delay in the shutoff of *Blimp-1*, *Hr3* and *crol* and delayed upregulation of *E74EF*, *E75B* and *E71CD*. When we investigated chromatin accessibility at these genes, we found that specific enhancers for Blimp-1 and Hr3 failed to close at 44h when cell cycle exit was disrupted while specific enhancers at E75B and E74EF failed to open (Fig. 6B,C). Our results suggest a model where ectopic E2F activity leads to delays in chromatin remodeling at specific ecdysone target genes, delaying their proper expression dynamics.

**Fig. 6.**
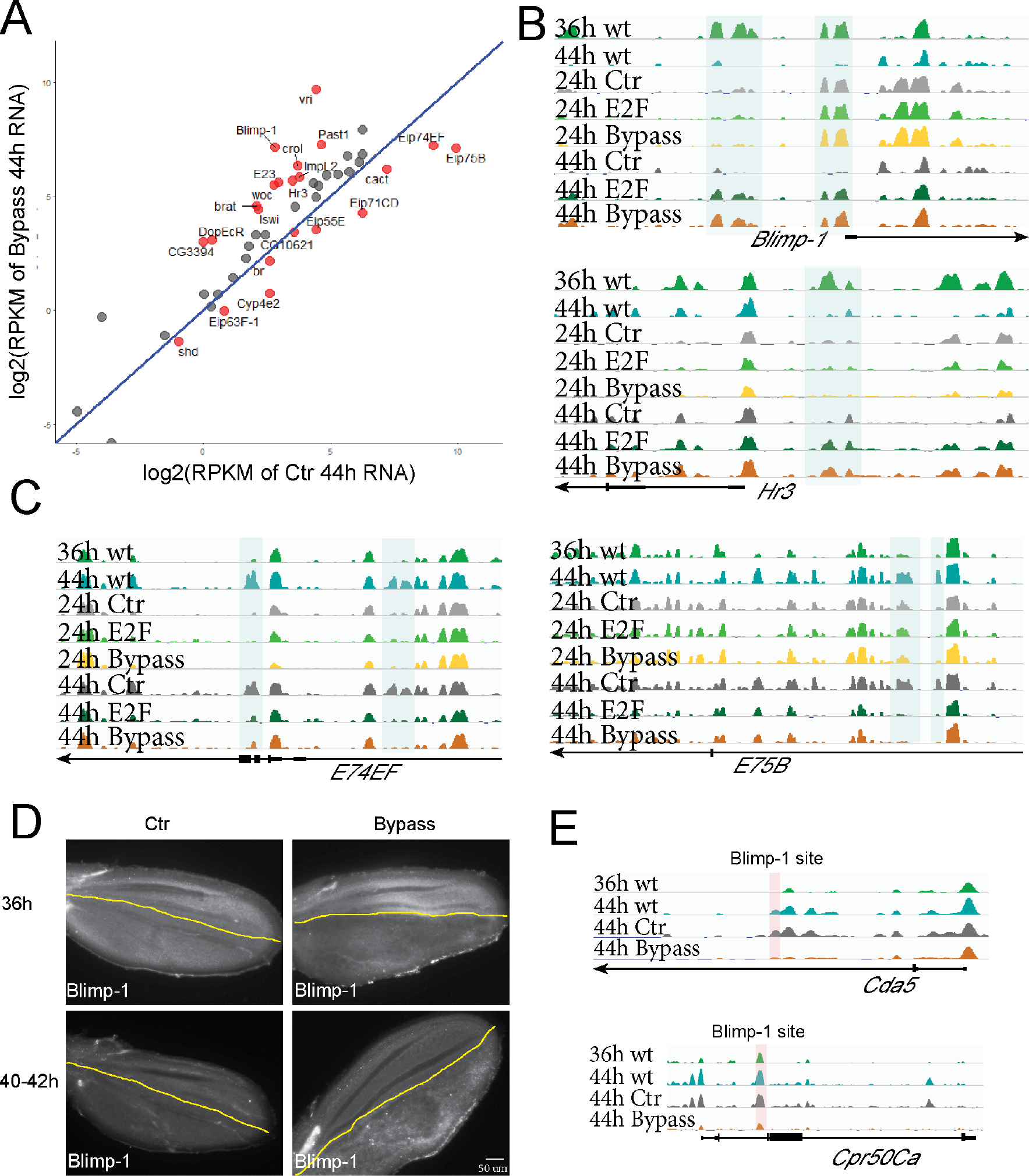
Bypassing cell cycle exit disrupts chromatin dynamics at ecdysone target genes and alters their expression. (A) Scatterplot of ecdysone responsive genes in 44h wings under conditions that bypass cell cycle exit vs. controls with normal exit. Genes with significant changes in expression are labeled in red. (B, C) Chromatin regions of *Blimp-1*, *Hr3*, *E74* and *E75* fail to close or open at 44h when cell cycle exit is compromised. (D) Blimp-1 antibody staining of wings at 36h and 40-42h wings with normal cell cycle exit (Ctr) or bypassed cell cycle exit in the posterior (using *engrailed*-Gal4/Gal80^ts^). Compromising cell cycle exit delays the activation of Blimp-1 in a compartment autonomous manner. (E) Peaks that fail to open at 44h from cuticle development genes harbor high scoring Blimp-1 binding sites.

We reasoned that these alterations in transcriptional regulators downstream of ecdysone signaling could lead to the alterations in chromatin accessibility for wing terminal differentiation genes when cell cycle exit is disrupted. Consistent with this, the Blimp-1 binding motif is significantly enriched in FAIRE peaks that are differentially regulated under conditions that delay or disrupt cell cycle exit (Supp. Fig. 10A). Several genes important for cuticle development such as *Cda5*, *Cpr50Ca*, *Cpr47Ec* and *TwdlT* harbor high scoring Blimp-1 binding sites and are likely direct Blimp-1 targets (Supp. Fig. 10B). Their peaks exhibit temporal dynamics consistent with a model where Blimp-1 either binds closed chromatin at 36h and facilitates subsequent chromatin opening at 44h or where high Blimp-1 binding at 36h somehow maintains closing that is lost when Blimp-1 levels plummet at 44h (Fig. 6E). The temporal and spatial resolution of our FAIRE timecourse is not sufficient to distinguish between these two scenarios. Interestingly, we also found a high scoring *Blimp-1* site in E74EF, suggesting its temporal regulation is also dependent on Blimp-1.

We considered the possibility that our genetic disruption of cell cycle exit could have non-autonomous effects that impact the timing or production of the ecdysone signal itself, leading to alterations in chromatin remodeling at specific targets. We therefore tested whether our manipulations of cell cycle exit impact Blimp-1 expression non-autonomously. For this, we expressed E2F+CycD specifically in the posterior compartment of the pupa wing using the *Engrailed*-Gal4/Gal80^ts^ system. Under these conditions only the posterior wing continues to proliferate while the anterior wing becomes postmitotic with the normal timing (Ma and Buttitta, 2017). We found that when we disrupted cell cycle exit in the posterior wing only, Blimp-1 protein levels were reduced at 36h but higher at 40-42h consistent with the delay in *Blimp-1* activation we observed by RNAseq. Importantly, Blimp-1 levels were unaffected in the anterior wing, showing the normal increase in Blimp-1 levels at 36h and drop in levels at 44h. This demonstrates that disrupting cell cycle exit impacts the timing of ecdysone target gene expression in a compartment-autonomous manner (Fig.6D), consistent with our findings on compartment-specific effects on cuticle formation (Fig. 5E). Our data are consistent with a model where the cis regulatory DNA at genes encoding hormone-regulated transcription factors such as Blimp-1 responds to ectopic cell cycles or E2F activity to coordinate cell cycle exit with later steps of terminal differentiation downstream of the hormone pulses. This in turn leads to delays in chromatin remodeling at their targets and altered expression dynamics of downstream wing terminal differentiation genes (Fig. 7)

**Fig. 7.**
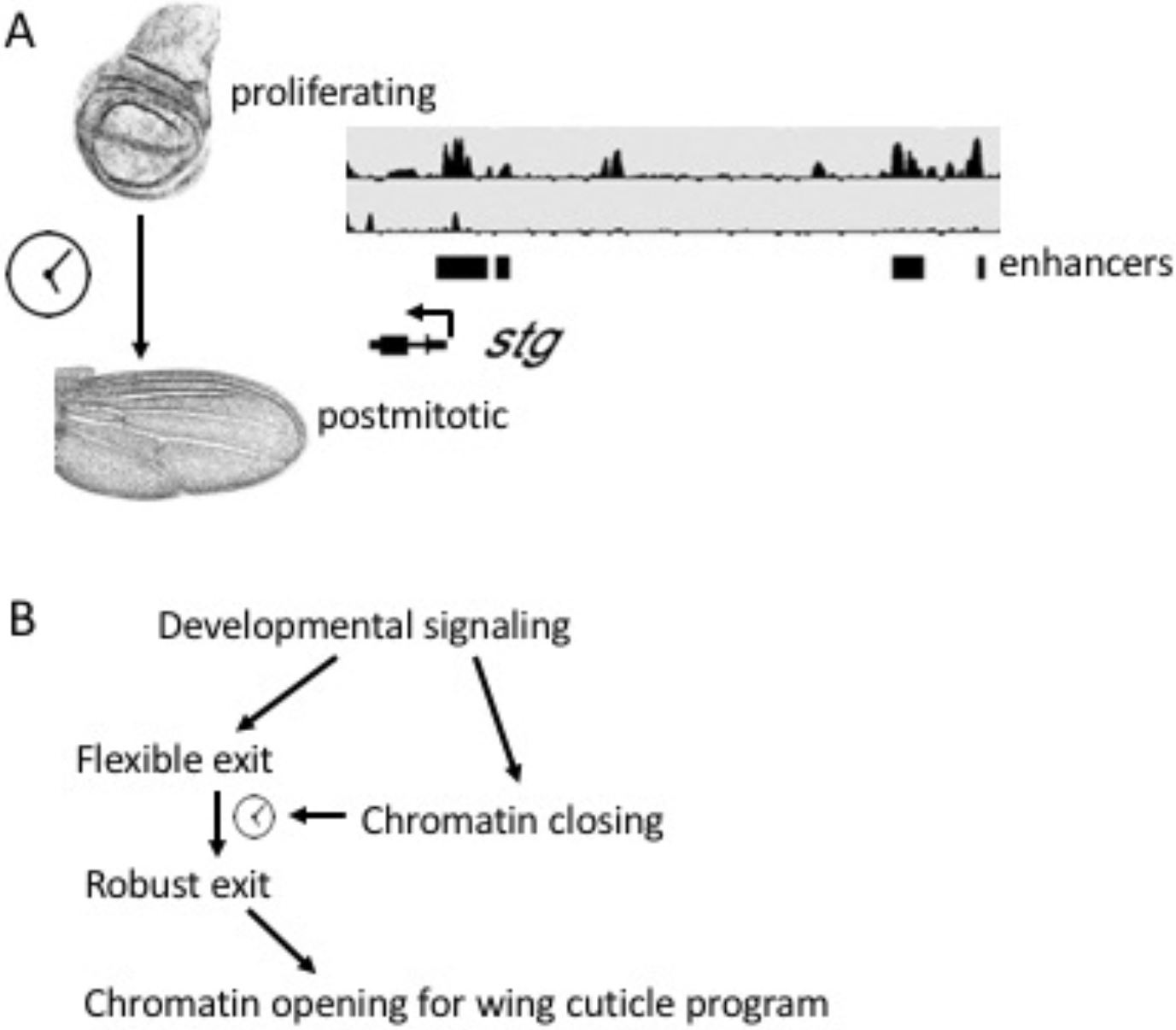
A model for the developmental coordination of cell cycle exit and chromatin accessibility. A. Regulatory elements at complex cell cycle genes such as *stg* become inaccessible in a developmentally controlled manner during Robust G0. This limits their activation in response to proliferative signals. B. While chromatin changes at cell cycle genes are developmentally controlled, delaying or disrupting cell cycle exit impacts the opening of chromatin at genes in the wing terminal differentiation program that are potentially controlled via transcription factors downstream of ecdysone signaling.

## Discussion

### Most chromatin and gene expression changes in the wing during metamorphosis are independent of cell cycling status

A striking feature of the *Drosophila* wing during metamorphosis is the coordination between the cell cycle and tissue morphogenetic changes. We hypothesized that the synchronous exit from the cell cycle may impact chromatin accessibility and lead to widespread gene expression changes to coordinate differentiation with cell cycle exit. However, this is not the case. When we compromise cell cycle exit we observe relatively few changes in chromatin accessibility. We therefore conclude that the majority of chromatin accessibility dynamics during metamorphosis are developmentally programmed and occur coincident with cell cycle exit, but are independent of the transition from a proliferating to a postmitotic state.

### Chromatin accessibility changes at cell cycle genes during metamorphosis

Terminal differentiation and the transition from proliferation to a postmitotic state are usually coupled during development. Cell cycle arrest has been proposed to be essential for terminal differentiation in several cell types by promoting or maintaining the proper expression of late differentiation genes (Matus et al., 2015; Nicolay et al., 2010; Novitch et al., 1999; Xu et al., 2014). While studies in other contexts have shown that cell cycle exit and overt terminal differentiation can be separable (Ajioka et al., 2007; Engerer et al., 2017; Korzelius et al., 2011; Mohamed et al., 2018; Sage et al., 2005), in this study we have comprehensively characterized the gene expression and gene regulatory mechanisms underlying these two processes by examining the transcriptome and open chromatin landscape changes during cell cycle exit. Our study reveals that during wing differentiation, chromatin accessibility and gene expression changes are temporally coordinated, and that distal regulatory elements at a subset of critical cell cycle genes (*cycE, stg, e2f1*) become inaccessible during terminal differentiation, even when cell cycle exit is compromised (Fig 3,4). We suggest that these changes in chromatin accessibility at essential and rate-limiting cell cycle genes provide additional barriers to cycling and provide a molecular explanation for the robust G0 state we observe after 36h APF. Notably, the closed distal regulatory elements at *cycE, stg* and *e2f1* in robust G0 contain known Su(H) and Yorkie binding sites (Supp. Fig. 8) (Djiane et al., 2013; Oh et al., 2013), in addition to predicted sites for transcription factors of pathways that promote proliferation in undifferentiated wings. Their closing likely explains why pupal wings after 36h fail to proliferate in response to many hyperplastic signalings, including direct activation of Notch or Yorkie (Buttitta et al., 2007).

Although the distal enhancers for *cycE* and *stg* remain inaccessible when we disrupt cell cycle exit (Fig.4), ectopic E2F + CycD can re-activate *cycE* and *stg* expression at 44h to support continued cycling (Fig. 4D). This is consistent with our previous finding that ectopic CycE + Stg is required in addition to E2F to keep cells in the wing cycling past robust G0 or to induce cell cycle re-entry from an established postmitotic G0 [7]. We suggest that ectopic E2F + CycD acts through accessible TSS proximal regulatory sites to re-activate *cycE* and *stg* expression.

The addition of ectopic CycD is essential for the reactivation of *stg* expression at 44h and continued cycling. Ectopic E2F alone does not reactivate *stg* nor support continued cycling, despite increasing the expression of many other cell cycle genes (Fig. 4). Why is adding CycD required for re-activation of *stg* expression? The most common model for CycD function is that it phosphorylates Rbf and weakens RB-mediated repression of E2F, thereby compromising DREAM repressive function. This would suggest that ectopic CycD may be needed to overcome additional dREAM repressive barriers at specific targets. However, recent work has suggested CycD activity may also play other roles to promote cell cycle entry from G0 (Narasimha et al., 2014). Further work to examine the regulatory elements that drive *cycE* and *stg* in this context and the proteins that bind them will be necessary.

Our model is consistent with recent findings in *C. elegans* and other contexts that chromatin remodeling plays an important role in ensuring cell cycle exit during differentiation (Albini et al., 2015; Ruijtenberg and van den Heuvel, 2015). However, by disrupting cell cycle exit rather than chromatin remodeling, we are able to largely uncouple terminal differentiation and cell cycle arrest in the fly wing. This allowed us to distinguish changes in chromatin accessibility at a subset of late differentiation genes that are dependent upon proper cell cycle arrest, from those throughout the majority of the genome, including cell cycle genes that are independent from arrest.

While we do not know the chromatin remodelers responsible for the closing of the distal regulatory elements at *cycE*, *stg* and *e2f1*, we do find conserved Ecdysone Receptor binding sites that exhibit peaks of EcR binding during pupal stages according to ModENCODE data (Negre et al., 2011). An attractive model is that the strong peak of ecdysone occurring at 24h triggers EcR complexes to recruit chromatin remodelers to subsequently modulate elements at differentiation genes and complex cell cycle genes to coordinate differentiation with cell cycle exit. As our disruptions of cell cycle exit do not seem to impact the pulse of ecdysone at 24h, closing of chromatin via this mechanism would proceed independent of cell cycling status and act to limit cell cycle entry even in the presence of strong ectopic cell cycle activation.

### Preventing cell cycle exit compromises a portion of the wing terminal differentiation program

Our results reveal that most chromatin accessibility changes at potential regulatory elements in fly wings are developmentally regulated and change independent of cell cycling status. However, compromising cell cycle exit does alter chromatin accessibility at a small subset of temporally regulated transcription factors that impact the proper timing of the wing cuticle differentiation program (Fig. 6F). We find that compromising cell cycle exit somehow cell-autonomously delays the temporal gene expression and chromatin remodeling cascade downstream of ecdysone signaling in the wing (Fig. 7). Our work implies an important role for Blimp-1 in the wing cuticle differentiation program. Blimp-1 is directly induced by ecdysone and is a transcriptional repressor that has been well studied for silencing *ftz-f1* at the onset of metamorphosis, as well as regulating cuticle formation in the fly embryo (Agawa et al., 2007; Akagi et al., 2016; Ozturk-Colak et al., 2016). Here we show that *Blimp-1* is also highly expressed at 36h APF wings, following the second and highest pulse of ecdysone during metamorphosis. *Blimp-1* is then immediately silenced by 44h. We found that dynamic chromatin regions that open at 44h are enriched for Blimp-1 binding sites. Some of these sites are potential regulatory elements for cuticle genes and other ecdysone targets such as *E74EF* (Supp. Fig. 10). Since these regions are closed when Blimp-1 is present and only open after Blimp-1 goes away, we propose that Blimp-1 blocks the accessibility of these dynamic regulatory elements. Consistent with this model, we also identified Blimp-1 binding sites at a dynamic open region at the *ftz-f1* locus. This region is transiently open at 6h when Blimp-1 is absent and *ftz-f1* is expressed, mirroring the pattern of accessibility for the potential Blimp-1 site at *E74EF* at 44h. Future work will focus on revealing the proximal factor that acts on chromatin accessibility at Blimp-1 regulatory elements to link cell cycling status with the temporal wing differentiation program.

## Materials and Methods

### Fly stocks and genetics

FAIRE and RNA seq samples with genetic manipulations:

*w/ y, w, hs-FLP; tub>CD2>GAL4, UAS-GFP/ +; tub-gal80TS/ +*
*w/ y, w, hs-FLP; tub>CD2>GAL4, UAS-GFP/ UAS-E2F1, UAS-DP; tub-gal80TS/ +*
*w/ y, w, hs-FLP; tub>CD2>GAL4, UAS-GFP/ UAS-CycD, UAS-Cdk4; tub-gal80TS/ UAS-E2F1, UAS-DP*

Transgenes are described in (Buttitta et al., 2007). The *tub>CD2>Gal4* is from (Pignoni and Zipursky, 1997) and UAS E2F1, DP, CycD, Cdk4 are from (Neufeld et al., 1998).

Crosses were set up at 25 °C. Second instar larva (L2) were heat shocked at 37 °C for 42 min, then kept at 18 °C. White prepupa were collected for staging and kept at 18 °C until the equivalent of 12h APF at 25°C. Then pupa were shifted to 28 °C until the equivalent of 24h APF or 44h APF at 25°C for dissection. Pupae develop 2.2 times more slowly at 18°C than at 25°C and 1.2 times faster at 28°C than 25°C. All timepoints were adjusted to equivalent stages at 25°C for figures.

TEM, Chitin and Phosphohistone H3 staining:

*w/y, w, hs-FLP; Ap-GAL4, UAS-GFP/ +; tub-gal80TS/ +*
*w/y, w, hs-FLP; Ap-GAL4, UAS-GFP/ UAS-E2F1, UAS-DP; tub-gal80TS/ +*
*w/y, w, hs-FLP; Ap-GAL4, UAS-GFP/ UAS-CycD, UAS-Cdk4; tub-gal80TS/ UAS-E2F1, UAS-DP*

Crosses were set up and kept at 18 °C. White prepupa were collected and aged to the equivalent of 12h APF, then shifted to 28 °C until the equivalent of 24h APF, 44h APF (for PH3 staining) or 64h APF (for TEM and chitin staining).

Blimp-1 antibody staining:

*w/y, w, hs-FLP; en-GAL4, UAS-GFP/ +; tub-gal80TS/ +*
*w/y, w, hs-FLP; en-GAL4, UAS-GFP/ UAS-CycD, UAS-Cdk4; tub-gal80TS/ UAS-E2F1, UAS-DP*
*w/y, w, hs-FLP; en-GAL4, UAS-GFP/ Blimp-1^RNAi^ (BL 57479), UAS-DP; tub-gal80TS/ +*
*w/y, w, hs-FLP; en-GAL4, UAS-GFP/ +; tub-gal80TS/ white^RNAi^*

Crosses were set up and kept at 18 °C. White prepupa were collected and shifted to 28 °C until the equivalent of 36h APF or 40-42h APF for immunostaining.

PCNA reporter assay:

*PCNA-EmGFP/ y, w, hs-FLP; +; act>CD2>gal4, UAS-RFP/+*
*PCNA-EmGFP/ y, w, hs-FLP; UAS-E2F1, UAS-DP /+; act>CD2>gal4, UAS-RFP/+*
*PCNA-EmGFP/ y, w, hs-FLP; +/UAS-CycD, UAS-Cdk4; act>CD2>gal4, UAS-RFP/ UAS-E2F1, UAS-DP*

The *PCNA-EmGFP* line is described in (Swanhart et al., 2007). Crosses were set up and kept at 25 °C. White prepupa were collected and incubated to 26h APF then heat shocked at 37 °C for 12 min and incubated at 25 °C until 42h APF for dissection.

Enhancer-Gal4 reporters:

Transgenic flies were crossed with UAS-GFP (*cpr51A* region, VT016704) or UAS-destabilized GFP (*stg* region, BL45586 and e2f1 region, VT045332) and incubated at 25 °C. Then larval or pupal samples (staged from white prepupae) were dissected and immuno-stained for GFP.

### Sample preparation and data analysis for high-throughput sequencing

FAIRE samples and RNA samples were prepared as described previously (Uyehara et al., 2017). FAIRE-seq sequencing reads were aligned to the dm6 reference genome using Bowtie2 (Langmead and Salzberg, 2012). FAIRE-seq peak calling were performed using MACS2 and PePr (Zhang et al., 2014; Zhang et al., 2008) with q-value threshold at 0.01, and only common peaks from both programs were utilized for further analysis. Z-scores were calculated using the mean and standard deviation per chromosome arm. High fidelity peaks were chosen from peaks with maximal Z-score larger than 2. FAIRE-seq line plots were generated using deepTools (Ramirez et al., 2016). FAIRE-seq were visualized using Integrative Genomics Viewer (Robinson et al., 2011). DNA-binding motifs used for enrichment analysis were obtained from FlyFactorSurvey (Zhu et al., 2011). Motif de novo discovery, comparison with known motif and motif enrichment analysis were using the MEME tool, TOMTOM tool and AME tool in MEME suite (Bailey et al., 2009). Annotation of FAIRE peaks were carried out by assigning peaks to nearest TSS in R package ChIPpeakAnno (Zhu et al., 2010). RNA-seq sequencing reads were aligned to the dm6 reference genome using STAR and further counted using HTSeq (Anders et al., 2015; Dobin et al., 2013). RPKM values of RNA-seq were calculated through Cufflinks (Trapnell et al., 2010). Differentially expressed genes were defined as those having RPKM >1 in at least one stage and changing by at least twofold between pairwise time points. GO analysis was performed using DAVID (Database for Annotation, Visualization, and Integrated Discovery) (Huang da et al., 2009). All the statistical comparisons are carried out in DEseq2 (Love et al., 2014).

### Immunostaining and Microscopy

Immunostaining procedures were carried out as previously described (Ma and Buttitta, 2017). Primary antibodies used in this study include: Anti-phospho-Ser10 histone H3, 1:2000 rabbit (Millipore #06-570) or mouse (Cell Signaling #9706); Anti-GFP, 1:1000 chicken (Life Technologies A10262) or 1:1000 rabbit (Life Technologies A11122); Anti-Blimp-1, 1:500 rabbit (Active motif 61054). DNA was labeled by 1 ug/ml DAPI in 1× PBS, 0.1% Triton X for 10 min and chitin was stained by 50 ug/ml Fluorescent Brightener 28 (Sigma-Aldrich, F3543) in 1× PBS, 0.1% Triton X for 10 min. Images were obtained using a Leica SP5 confocal (chitin staining) or Leica DMI6000B epifluorescence system.

### Transmission electron microscopy

Tissue was incubated in Karnovsky’s fixative for at least 1hr at room temperature, then overnight at 4 degrees. Samples were washed with 20x volume Sorenson’s buffer 3x, before post-fixing in 2% osmium tetroxide in Sorenson’s buffer for 1hr at RT. Tissue was again washed 3x with 20x volume Sorenson’s buffer, then dehydrated through ascending concentrations of acetone and embedded in EMbed 812 epoxy resin. Semi-thin sections were stained with toluidine blue for tissue identification. Selected regions of interest were ultra-thin sectioned 70 nm in thickness and post stained with uranyl acetate and Reynolds lead citrate. They were examined using a JEOL JEM-1400 Plus transmission electron microscope (TEM) at 80 kV.

## Acknowledgements

We thank Ajai Pulianmackal, Shyama Nandakumar, Kerry Flegel and Dan Sun for their help in preparing FAIRE-seq and RNA-seq samples. We thank Emily Rozich for help with examining he CyclinE-GFP reporter. We thank Spencer Nystrom for help with bioinformatic analysis. We thank Pennelope Blakely (Microscopy & Image-analysis Lab, Biomedical Research Core Facilities, University of Michigan) and Stephen Ireland for the help with TEM experiments. The authors have no conflicts of interest.

## Acknowledgements

This work was supported by the American Cancer Society (RSG-15-161-01-DDC to LAB) and the NIH NIGMS (GM127367 to LAB).

## Supplemental Figures and Legends

**Supplemental Fig. 1.**
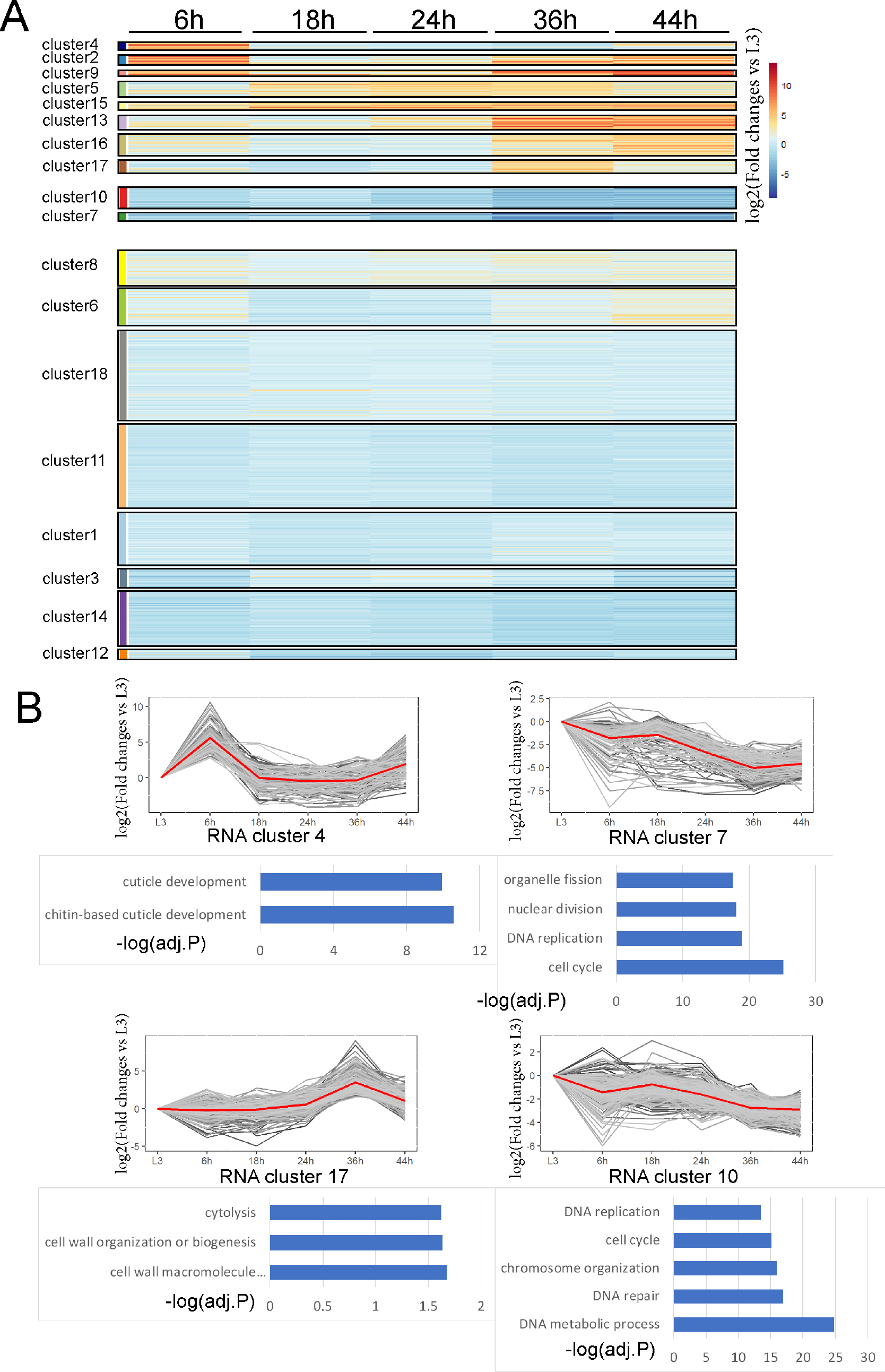
Gene expression is dynamic during metamorphosis. (A) Heatmap of RNA log2 fold change vs L3 for the indicated stages. The pattern of RNA changes during metamorphosis is separated into 18 k-means clusters. (B) Line plots of the log2 fold change vs L3 for the indicated RNA clusters. Each gene is represented by a single gray line and the average of all genes for the given cluster is plotted in red line. GO term enrichments are also shown along with their adjust P-values. During metamorphosis differentiation related genes such as cuticle development are activated while cell cycle genes are repressed.

**Supplemental Fig. 2.**
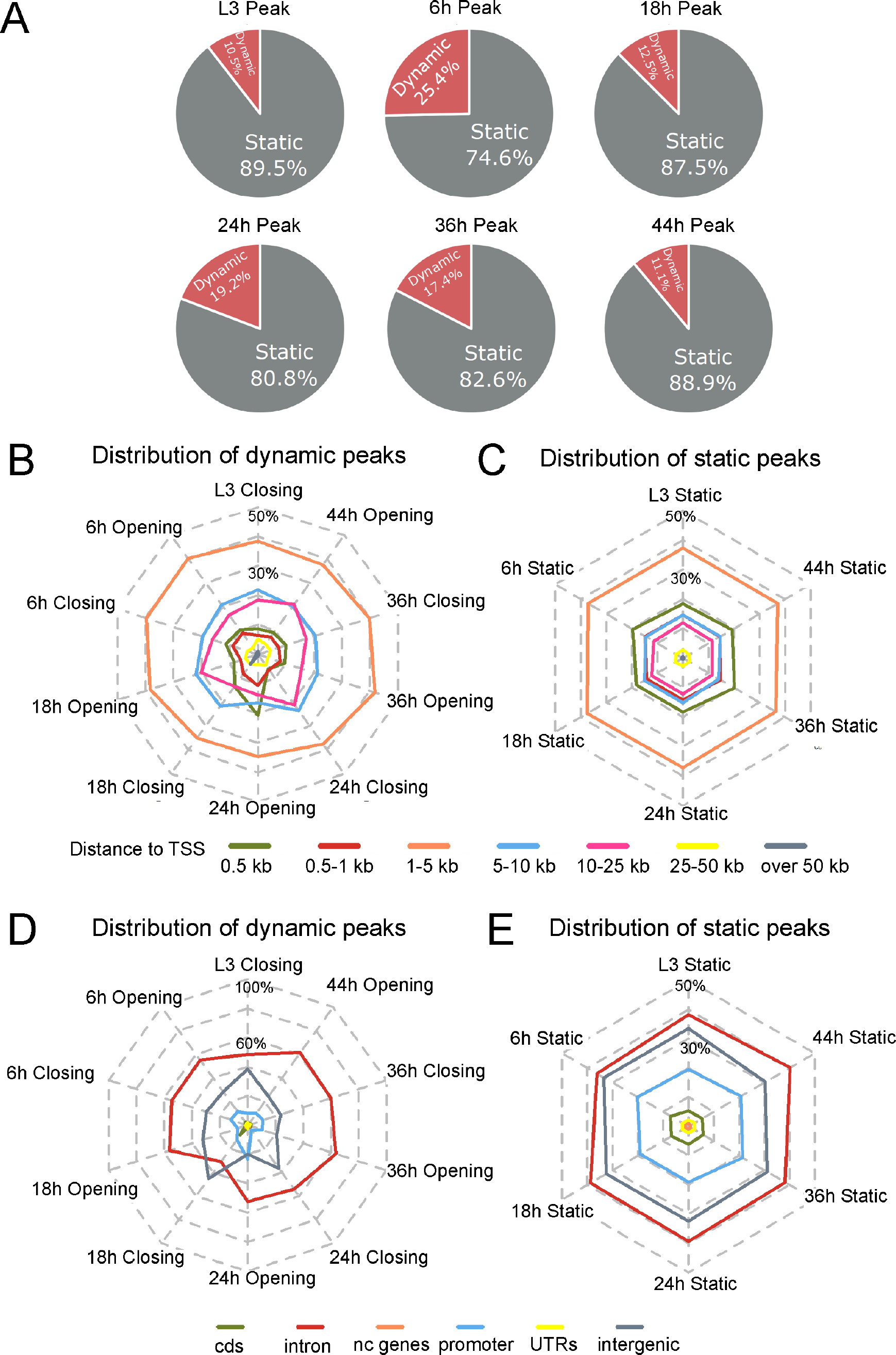
Locations of dynamic vs. static open chromatin. (A) Pie charts of the proportion of dynamic peaks and static peaks of each stage examined. Peaks without significant changes (<2-fold) between neighboring timepoints were defined as “static”. Peaks bearing changes by >2-fold were defined as “dynamic”. (B, C) Radar charts display the distribution of indicated dynamic (B) and static (C) peak categories in different distances to TSS. (D, E) Radar charts display the distribution of indicated dynamic (D) and static (E) peak categories in cds, intron, non-coding genes (nc genes), proximal promoter (−500bp to 150 bp of TSS), UTRs and intergenic regions. For dynamic peaks, “closing” is defined as peaks that decrease in accessibility by >2-fold comparing to the previous stage, conversely “opening” indicates peaks that increase in accessibility by >2-fold comparing to the previous stage.

**Supplemental Fig. 3.**
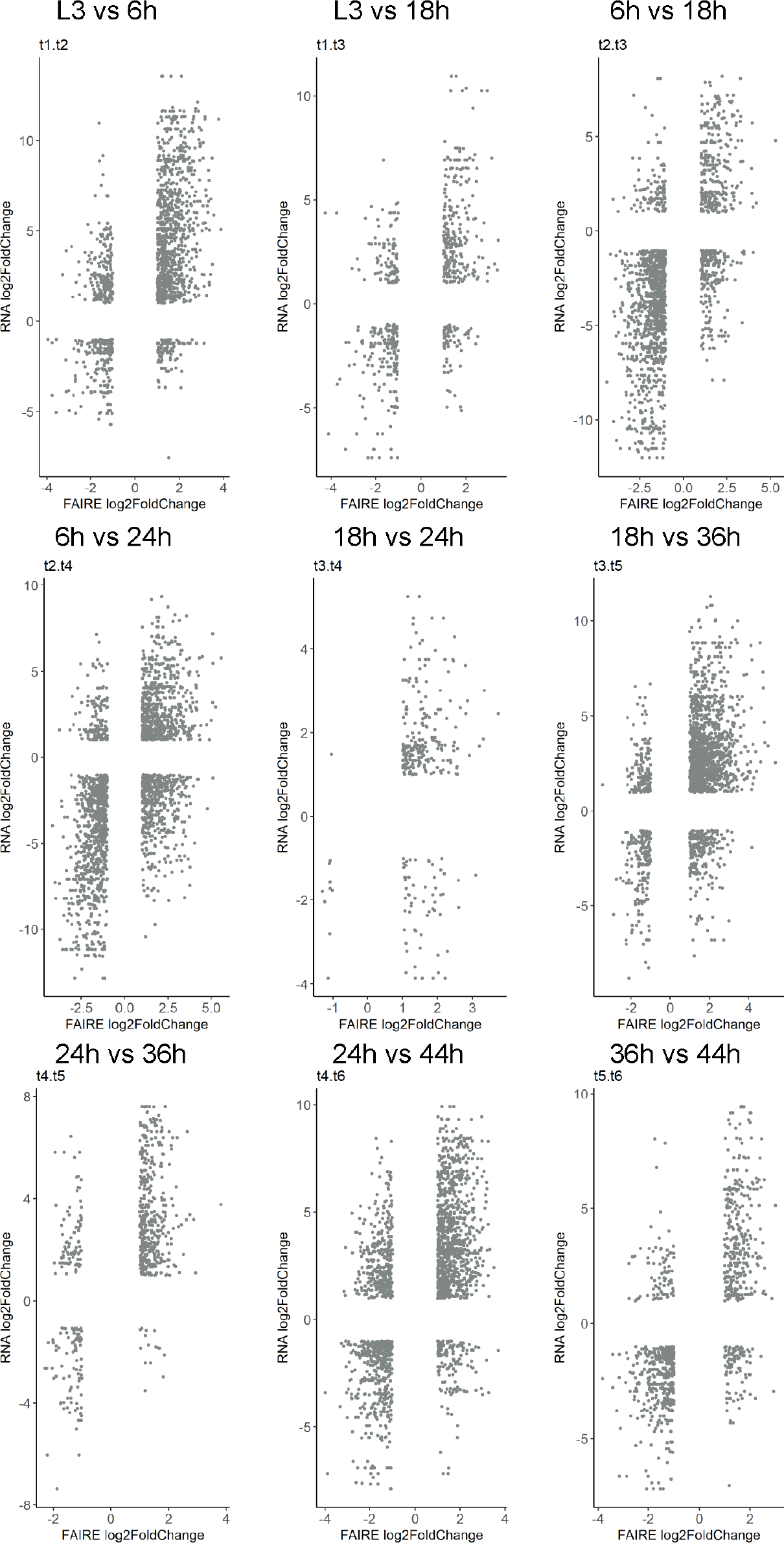
The majority of dynamic open chromatin is associated with gene activation rather than gene repression. Scatterplots of FAIRE peaks and corresponding genes with significant changes between 2 sequential stages. Significance is defined by 2-fold changes and adjust P-value less than 0.05.

**Supplemental Fig. 4.**
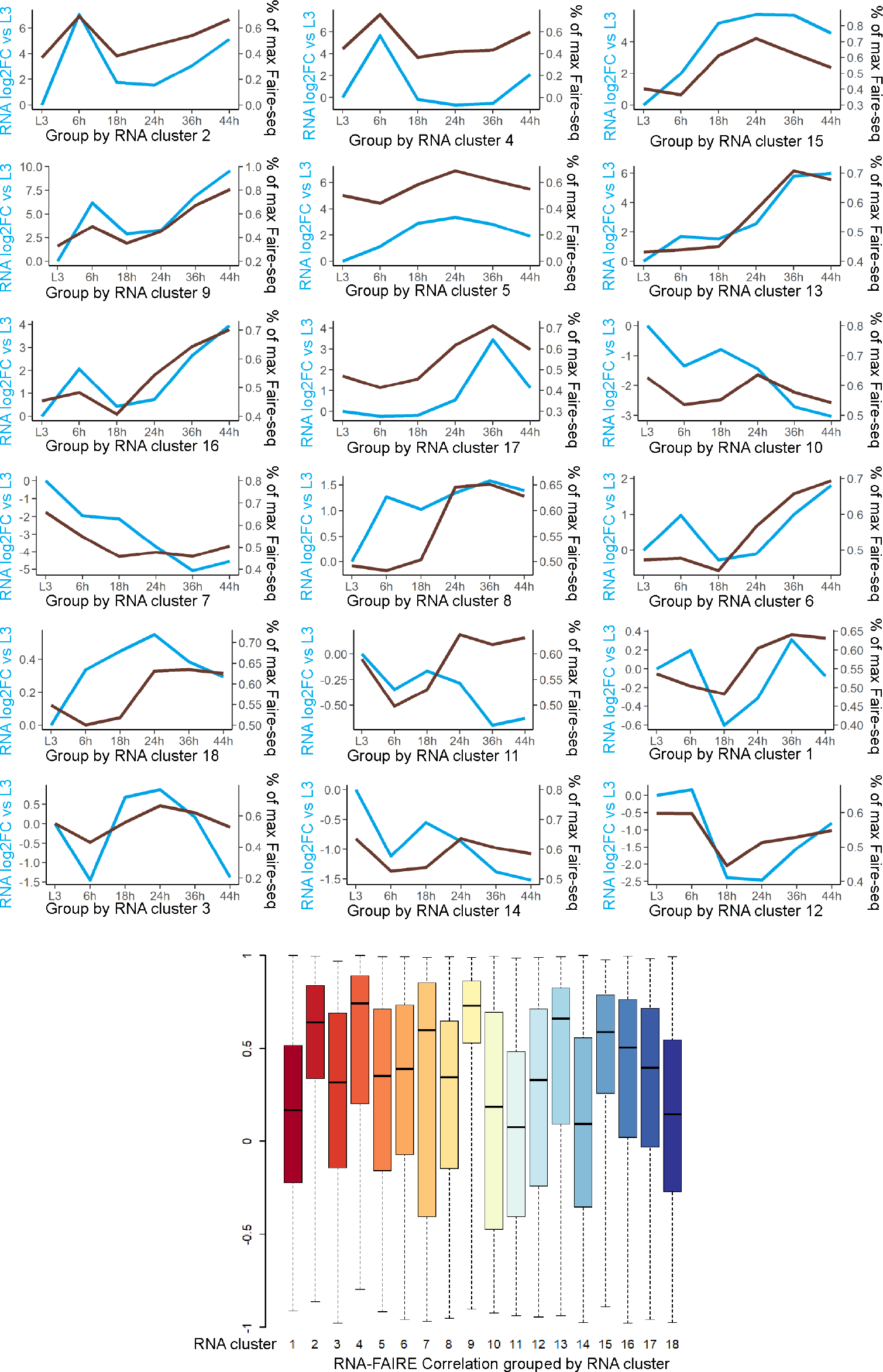
Coordination of RNA and FAIRE peak changes grouped by RNA clustering. Trajectories of average changes between genes and their corresponding FAIRE peaks over the 6 stages for each of the 18 RNA clusters. Boxplot of the Pearson correlation coefficients between RNA and FAIRE for each RNA cluster is shown.

**Supplemental Fig. 5.**
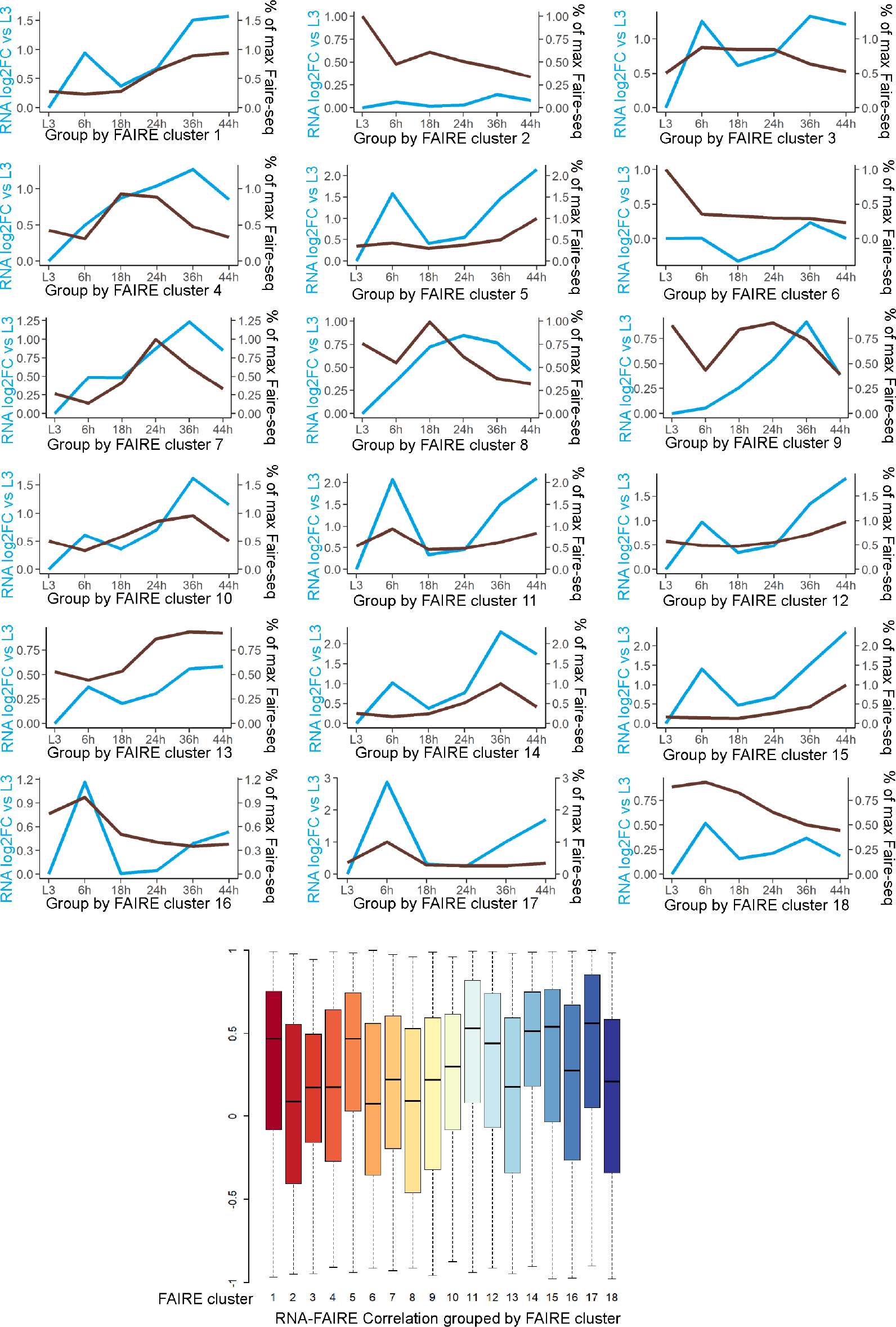
Coordination of RNA and FAIRE peak changes grouped by FAIRE peak clustering. Trajectories of average changes between FAIRE peaks and their corresponding genes over the 6 stages for each of the 18 FAIRE clusters. Boxplot of the Pearson correlation coefficients between RNA and FAIRE for each FAIRE cluster is shown.

**Supplemental Fig. 6.**
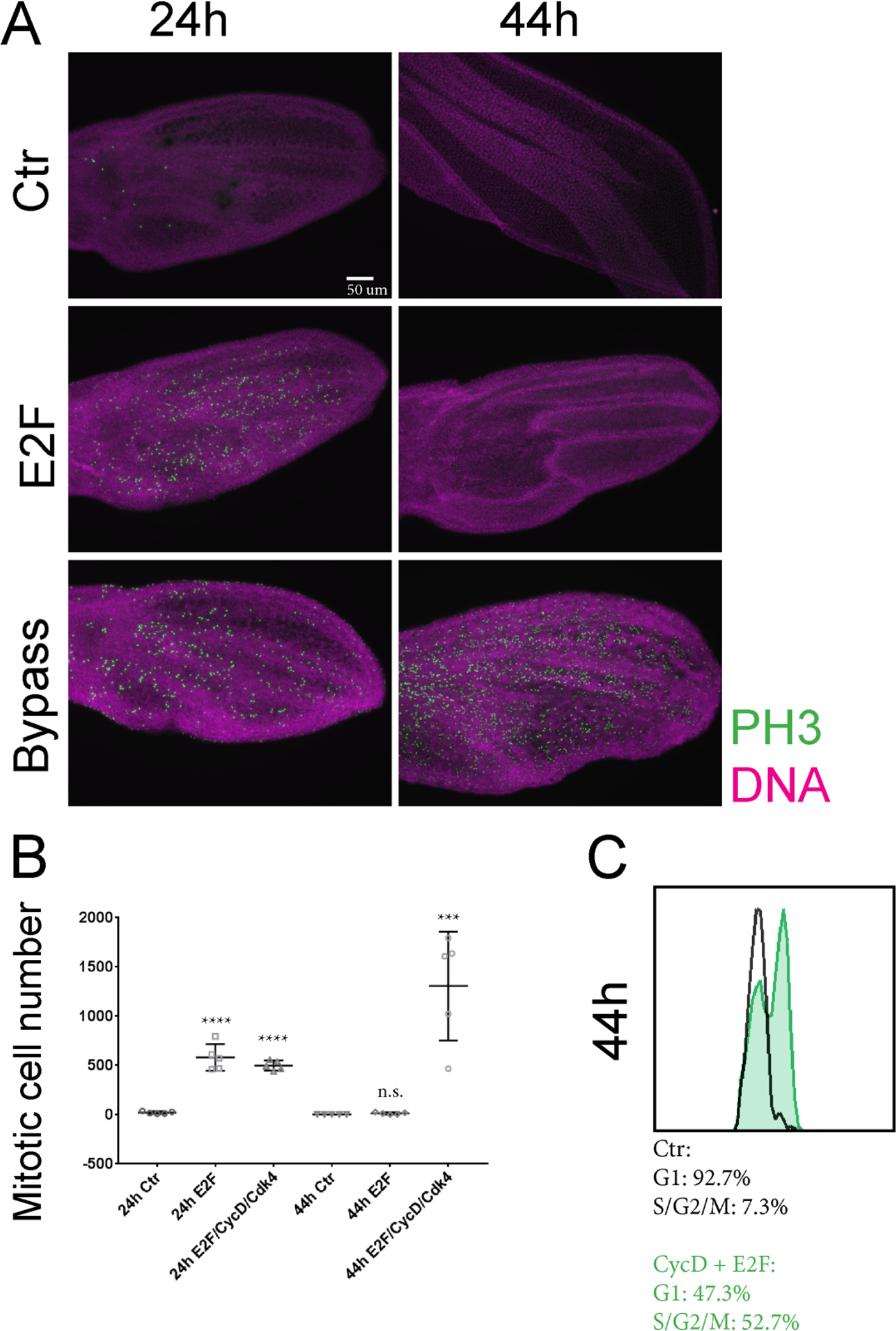
Two stages of G0 in differentiating wings. (A) E2F or E2F/CycD/Cdk4 (bypass) was overexpressed in the dorsal layer of wing epithelia under the control of *Apterous*-Gal4/Gal80^ts^ from 12h APF. 24h and 44h wings were immunostained against phospho-histone H3 (ph3). (B) The number of PH3 spots of each wing is counted and 5 wings for each genotype are quantified. (C) Cell cycle profile of the FAIRE samples that bypassed robust G0 by E2F/CycD/Cdk4 was examined by FACS. P-values were determined by an unpaired t-test; **** <0.0001, ***<0.001.

**Supplemental Fig. 7.**
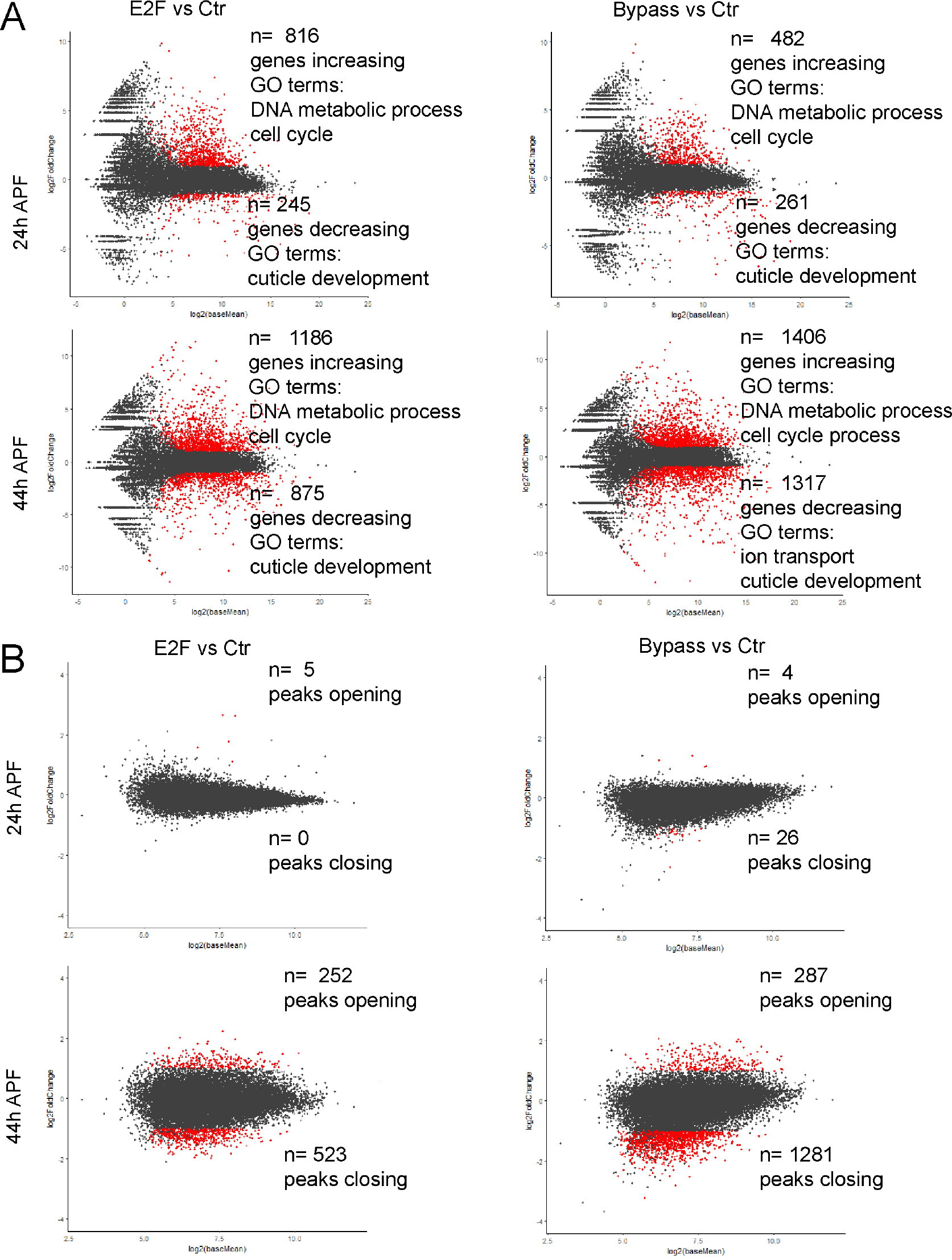
RNA-seq and FAIRE-seq changes when G0 is delayed (E2F expression wings) or bypassed (E2F/CycD/Cdk4 expression wings) MA-plots of RNA (A) and FAIRE (B) changes of 24 and 44h wings compared to control. Genes and peaks that are significant in changes with 2-fold difference and adjusted P-value less than 0.05 are labeled in red.

**Supplemental Fig. 8.**
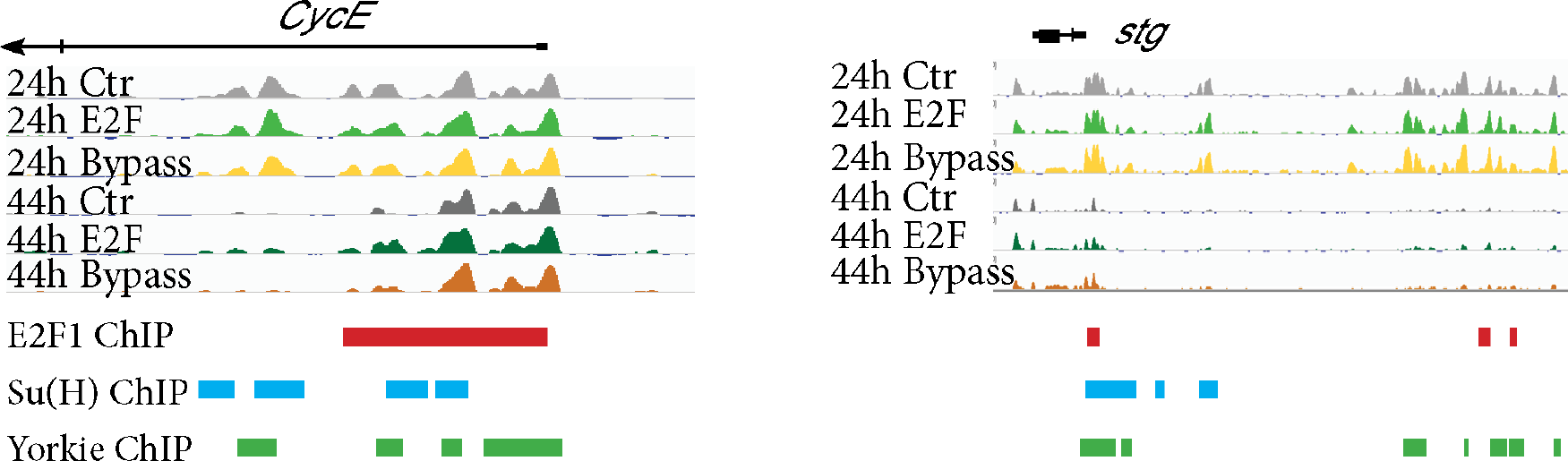
Locations of transcription factor binding sites in *cycE*, *stg* and *e2f1*. TSS and dynamic accessible chromatin regions of *cycE* and *stg* contain E2F1 (Korenjak et al., 2012), Su(H) (Djiane et al., 2013) and Yorkie/Sd (Oh et al., 2013) binding sites.

**Supplemental Fig. 9.**
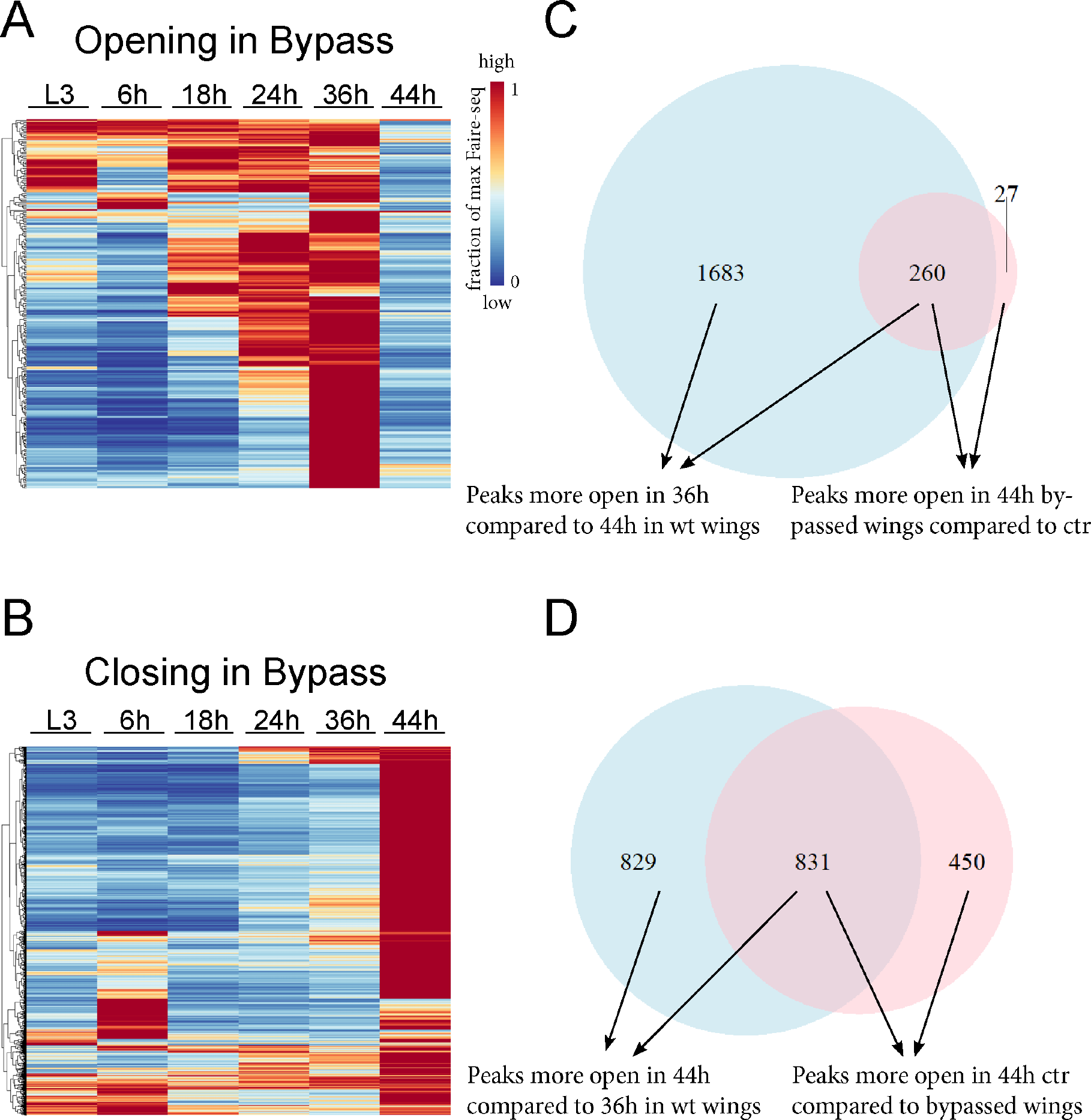
Bypassing cell cycle exit disrupts the temporal dynamics of chromatin accessibility at a subset of genes. (A, B) Heatmap shows the temporal dynamics during normal development for the peaks that are more accessible or less accessible at 44h wings expressing E2F/CycD/Cdk4, plotted as a fraction of the maximum FAIRE rpkm value. Compromising G0 leads to the failure of proper closing of 36h peaks as well as the opening of 44h peaks. (C) Overlap between peaks that normally open at 36h in wt and peaks more accessible at 44h bypassed wings. (D) Overlap between peaks that normally open at 44h in wt and peaks less accessible at 44h bypassed wings.

**Supplemental Fig. 10.**
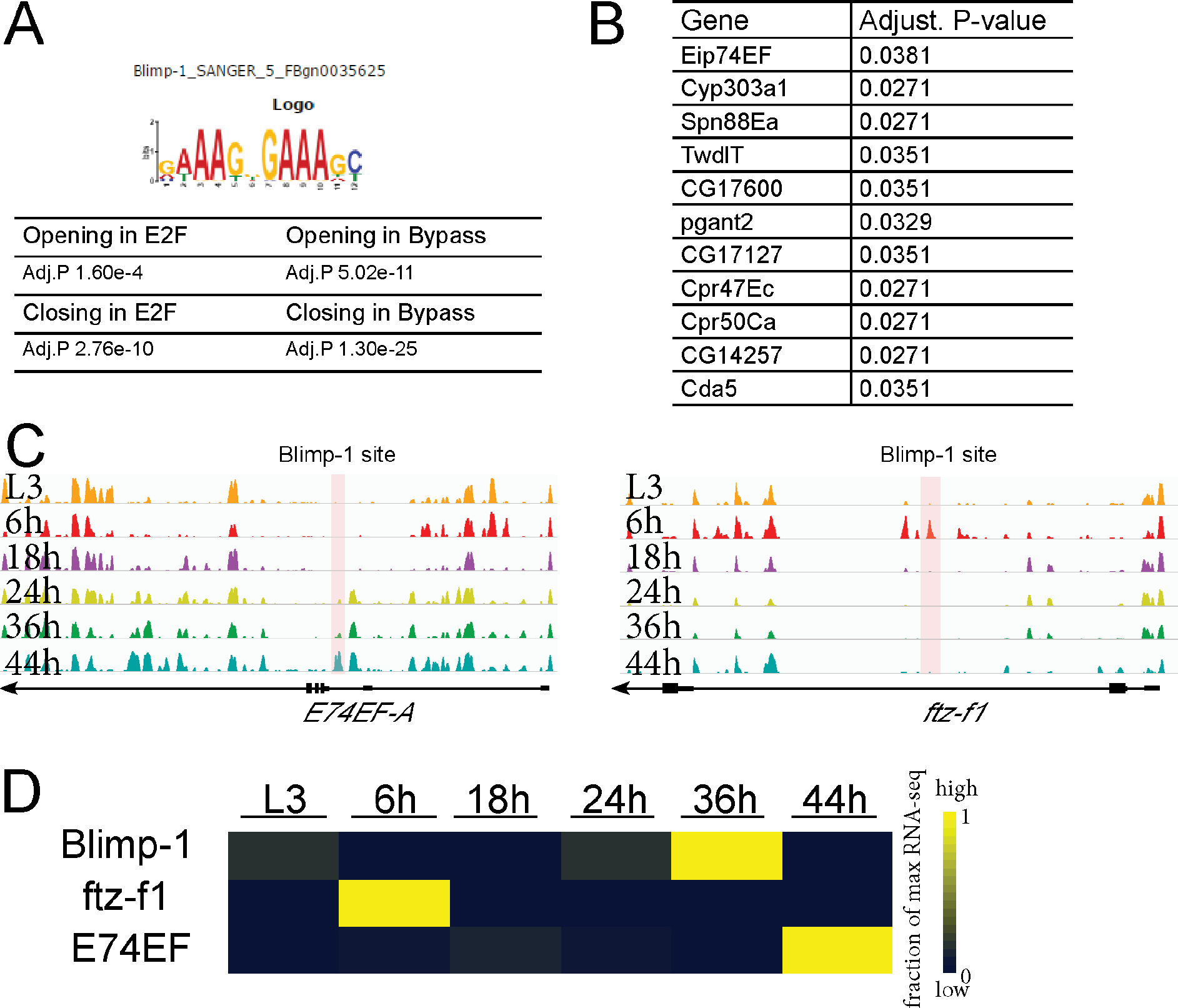
Compromising G0 disrupts the temporal dynamics of potential Blimp-1 targets. (A) Blimp-1 motif is enriched in the dynamic peaks disrupted by E2F or bypass through AME analysis. (B) List of genes containing peaks that fail to open at 44h with high scoring Blimp-1 binding sites. (C) Chromatin accessibility changes at *E74EF* and *ftz-f1* locus with Blimp-2 binding sites shown. (D) Expression changes of *Blimp-1, ftz-f1* and *E74EF* during normal development.

**Supplemental Fig. 11.**
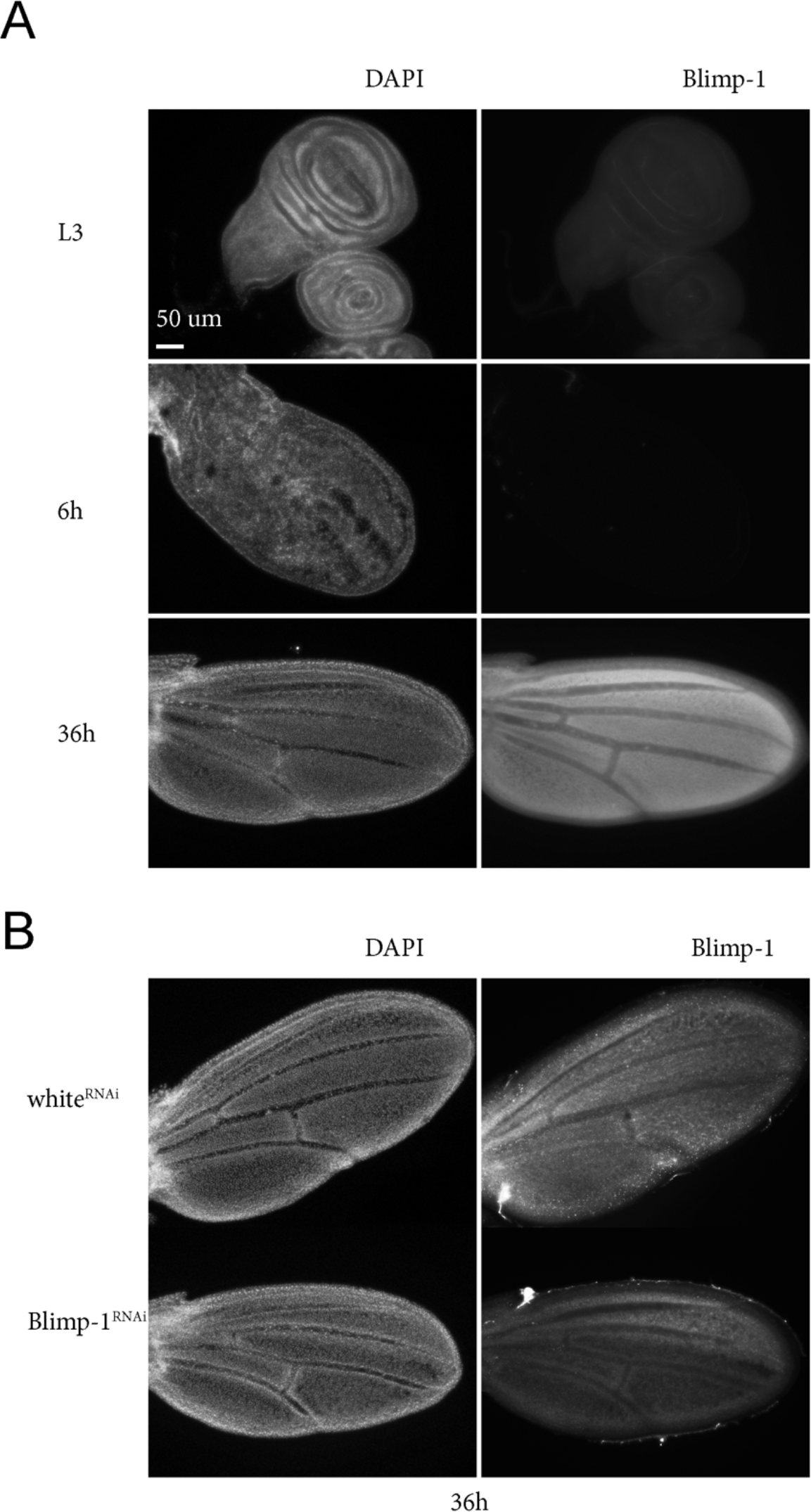
Validation of Blimp-1 reagents. (A) Blimp-1 antibody staining in wildtype L3, 6h and 36h wings corresponds to the gene expression changes of *Blimp-1*. (B) Expressing Blimp-^RNAi^ in the posterior wings by *engrailed*-Gal4/Gal80^ts^ from 0h APF reduces the level of Blimp-1 protein at 36h wings.

## References

1. O’Farrell PH. Quiescence: early evolutionary origins and universality do not imply uniformity. Philos Trans R Soc Lond B Biol Sci. 2011;366:3498–507. doi:10.1098/rstb.2011.0079.

2. Soufi A, Dalton S. Cycling through developmental decisions: how cell cycle dynamics control pluripotency, differentiation and reprogramming. Development. 2016;143:4301–11. doi:10.1242/dev.142075.

3. Ruijtenberg S, van den Heuvel S. Coordinating cell proliferation and differentiation: Antagonism between cell cycle regulators and cell type-specific gene expression. Cell Cycle. 2016;15:196–212. doi:10.1080/15384101.2015.1120925.

4. Sadasivam S, DeCaprio JA. The DREAM complex: master coordinator of cell cycle-dependent gene expression. Nat Rev Cancer. 2013;13:585–95. doi:10.1038/nrc3556.

5. van den Heuvel S, Dyson NJ. Conserved functions of the pRB and E2F families. Nat Rev Mol Cell Biol. 2008;9:713–24. doi:10.1038/nrm2469.

6. Blais A, Dynlacht BD. E2F-associated chromatin modifiers and cell cycle control. Curr Opin Cell Biol. 2007;19:658–62. doi:10.1016/j.ceb.2007.10.003.

7. Buttitta LA, Katzaroff AJ, Perez CL, de la Cruz A, Edgar BA. A double-assurance mechanism controls cell cycle exit upon terminal differentiation in Drosophila. Dev Cell. 2007;12:631–43. doi:10.1016/j.devcel.2007.02.020.

8. Firth LC, Baker NE. Extracellular signals responsible for spatially regulated proliferation in the differentiating Drosophila eye. Dev Cell. 2005;8:541–51. doi:10.1016/j.devcel.2005.01.017.

9. The I, Ruijtenberg S, Bouchet BP, Cristobal A, Prinsen MBWW, Van Mourik T, et al. Rb and FZR1/Cdh1 determine CDK4/6-cyclin D requirement in C. elegans and human cancer cells. Nat Commun. 2015;6:5906. doi:10.1038/ncomms6906.

10. Buttitta LA, Katzaroff AJ, Edgar BA. A robust cell cycle control mechanism limits E2F-induced proliferation of terminally differentiated cells in vivo. J Cell Biol. 2010;189:981–96. doi:10.1083/jcb.200910006.

11. Camarda G, Siepi F, Pajalunga D, Bernardini C, Rossi R, Montecucco A, et al. A pRb-independent mechanism preserves the postmitotic state in terminally differentiated skeletal muscle cells. J Cell Biol. 2004;167:417–23. doi:10.1083/jcb.200408164.

12. Pajalunga D, Tognozzi D, Tiainen M, D’Angelo M, Ferrantelli F, Helin K, et al. E2F activates late-G1 events but cannot replace E1A in inducing S phase in terminally differentiated skeletal muscle cells. Oncogene. 1999;18:5054–62. doi:10.1038/sj.onc.1202897.

13. Cecchini MJ, Thwaites MJ, Talluri S, MacDonald JI, Passos DT, Chong J-L, et al. A retinoblastoma allele that is mutated at its common E2F interaction site inhibits cell proliferation in gene-targeted mice. Mol Cell Biol. 2014;34:2029–45. doi:10.1128/MCB.01589-13.

14. Korzelius J, The I, Ruijtenberg S, Prinsen MBW, Portegijs V, Middelkoop TC, et al. Caenorhabditis elegans cyclin D/CDK4 and cyclin E/CDK2 induce distinct cell cycle re-entry programs in differentiated muscle cells. PLoS Genet. 2011;7:e1002362. doi:10.1371/journal.pgen.1002362.

15. Ebelt H, Zhang Y, Kampke A, Xu J, Schlitt A, Buerke M, et al. E2F2 expression induces proliferation of terminally differentiated cardiomyocytes in vivo. Cardiovasc Res. 2008;80:219–26. doi:10.1093/cvr/cvn194.

16. van Rijnberk LM, van der Horst SEM, van den Heuvel S, Ruijtenberg S. A dual transcriptional reporter and CDK-activity sensor marks cell cycle entry and progression in C. elegans. PLoS One. 2017;12:e0171600. doi:10.1371/journal.pone.0171600.

17. Di Stefano V, Giacca M, Capogrossi MC, Crescenzi M, Martelli F. Knockdown of cyclin-dependent kinase inhibitors induces cardiomyocyte re-entry in the cell cycle. J Biol Chem. 2011;286:8644–54. doi:10.1074/jbc.M110.184549.

18. Yao G. Modelling mammalian cellular quiescence. Interface Focus. 2014;4:20130074–20130074. doi:10.1098/rsfs.2013.0074.

19. Blais A, van Oevelen CJCC, Margueron R, Acosta-Alvear D, Dynlacht BD. Retinoblastoma tumor suppressor protein-dependent methylation of histone H3 lysine 27 is associated with irreversible cell cycle exit. J Cell Biol. 2007;179:1399–412. doi:10.1083/jcb.200705051.

20. Sdek P, Zhao P, Wang Y, Huang C-J, Ko CY, Butler PC, et al. Rb and p130 control cell cycle gene silencing to maintain the postmitotic phenotype in cardiac myocytes. J Cell Biol. 2011;194:407–23. doi:10.1083/jcb.201012049.

21. Peric-Hupkes D, Meuleman W, Pagie L, Bruggeman SWM, Solovei I, Brugman W, et al. Molecular Maps of the Reorganization of Genome-Nuclear Lamina Interactions during Differentiation. Mol Cell. 2010;38:603–13. doi:10.1016/j.molcel.2010.03.016.

22. Ma Y, Buttitta L. Chromatin organization changes during the establishment and maintenance of the postmitotic state. Epigenetics Chromatin. 2017;10:53. doi:10.1186/s13072-017-0159-8.

23. Guo Y, Flegel K, Kumar J, McKay DJ, Buttitta LA. Ecdysone signaling induces two phases of cell cycle exit in Drosophila cells. Biol Open. 2016;5:1648–61. doi:10.1242/bio.017525.

24. Uyehara CM, Nystrom SL, Niederhuber MJ, Leatham-Jensen M, Ma Y, Buttitta LA, et al. Hormone-dependent control of developmental timing through regulation of chromatin accessibility. Genes Dev. 2017;31:862–75. doi:10.1101/gad.298182.117.

25. Johnson SA, Milner MJ. Cuticle secretion in Drosophila wing imaginal discs in vitro: parameters of exposure to 20-hydroxy ecdysone. Int J Dev Biol. 1990;34:299–307. http://www.ncbi.nlm.nih.gov/pubmed/2117462. Accessed 16 Mar 2018.

26. Halme A, Cheng M, Hariharan IK. Retinoids regulate a developmental checkpoint for tissue regeneration in Drosophila. Curr Biol. 2010;20:458–63. doi:10.1016/j.cub.2010.01.038.

27. Schubiger M, Carré C, Antoniewski C, Truman JW. Ligand-dependent de-repression via EcR/USP acts as a gate to coordinate the differentiation of sensory neurons in the Drosophila wing. Development. 2005;132:5239–48. doi:10.1242/dev.02093.

28. Ashburner M. Puffs, genes, and hormones revisited. Cell. 1990;61:1–3. http://www.ncbi.nlm.nih.gov/pubmed/2180581. Accessed 16 Mar 2018.

29. Thummel CS. Ecdysone-regulated puff genes 2000. Insect Biochem Mol Biol. 2002;32:113–20. http://www.ncbi.nlm.nih.gov/pubmed/11755052. Accessed 16 Mar 2018.

30. King-Jones K, Thummel CS. Nuclear receptors--a perspective from Drosophila. Nat Rev Genet. 2005;6:311–23. doi:10.1038/nrg1581.

31. Stoiber M, Celniker S, Cherbas L, Brown B, Cherbas P. Diverse Hormone Response Networks in 41 Independent Drosophila Cell Lines. G3 (Bethesda). 2016;6:683–94. doi:10.1534/g3.115.023366.

32. Fristrom D, Liebrich W. The hormonal coordination of cuticulin deposition and morphogenesis in Drosophila imaginal discs in vivo and in vitro. Dev Biol. 1986;114:1–11. http://www.ncbi.nlm.nih.gov/pubmed/3082697. Accessed 16 Mar 2018.

33. Sobala LF, Adler PN. The Gene Expression Program for the Formation of Wing Cuticle in Drosophila. PLoS Genet. 2016;12:e1006100. doi:10.1371/journal.pgen.1006100.

34. Taylor J, Adler PN. Cell rearrangement and cell division during the tissue level morphogenesis of evaginating Drosophila imaginal discs. Dev Biol. 2008;313:739–51. doi:10.1016/j.ydbio.2007.11.009.

35. Sotillos S, De Celis JF. Interactions between the Notch, EGFR, and decapentaplegic signaling pathways regulate vein differentiation during Drosophila pupal wing development. Dev Dyn. 2005;232:738–52. doi:10.1002/dvdy.20270.

36. O’Keefe DD, Thomas SR, Bolin K, Griggs E, Edgar BA, Buttitta LA, et al. Combinatorial control of temporal gene expression in the Drosophila wing by enhancers and core promoters. BMC Genomics. 2012;13:498. doi:10.1186/1471-2164-13-498.

37. Song L, Zhang Z, Grasfeder LL, Boyle AP, Giresi PG, Lee B-K, et al. Open chromatin defined by DNaseI and FAIRE identifies regulatory elements that shape cell-type identity. Genome Res. 2011;21:1757–67. doi:10.1101/gr.121541.111.

38. Thomas S, Li X-Y, Sabo PJ, Sandstrom R, Thurman RE, Canfield TK, et al. Dynamic reprogramming of chromatin accessibility during Drosophila embryo development. Genome Biol. 2011;12:R43. doi:10.1186/gb-2011-12-5-r43.

39. Arnold CD, Gerlach D, Stelzer C, Boryń ŁM, Rath M, Stark A. Genome-wide quantitative enhancer activity maps identified by STARR-seq. Science. 2013;339:1074–7. doi:10.1126/science.1232542.

40. Zabidi MA, Arnold CD, Schernhuber K, Pagani M, Rath M, Frank O, et al. Enhancer-core-promoter specificity separates developmental and housekeeping gene regulation. Nature. 2015;518:556–9. doi:10.1038/nature13994.

41. FitzGerald PC, Sturgill D, Shyakhtenko A, Oliver B, Vinson C. Comparative genomics of Drosophila and human core promoters. Genome Biol. 2006;7:R53. doi:10.1186/gb-2006-7-7-r53.

42. Ohler U, Liao G, Niemann H, Rubin GM. Computational analysis of core promoters in the Drosophila genome. Genome Biol. 2002;3:RESEARCH0087. http://www.ncbi.nlm.nih.gov/pubmed/12537576. Accessed 16 Mar 2018.

43. Gurudatta B V, Yang J, Van Bortle K, Donlin-Asp PG, Corces VG. Dynamic changes in the genomic localization of DNA replication-related element binding factor during the cell cycle. Cell Cycle. 2013;12:1605–15. doi:10.4161/cc.24742.

44. Williams LH, Fromm G, Gokey NG, Henriques T, Muse GW, Burkholder A, et al. Pausing of RNA polymerase II regulates mammalian developmental potential through control of signaling networks. Mol Cell. 2015;58:311–22. doi:10.1016/j.molcel.2015.02.003.

45. Swanhart LM, Sanders AN, Duronio RJ. Normal regulation of Rbf1/E2f1 target genes in Drosophila type 1 protein phosphatase mutants. Dev Dyn. 2007;236:2567–77. doi:10.1002/dvdy.21265.

46. Lehman DA, Patterson B, Johnston LA, Balzer T, Britton JS, Saint R, et al. Cis-regulatory elements of the mitotic regulator, string/Cdc25. Development. 1999;126:1793–803. http://www.ncbi.nlm.nih.gov/pubmed/10101114. Accessed 16 Mar 2018.

47. Richardson HE, O’Keefe L V, Reed SI, Saint R. A Drosophila G1-specific cyclin E homolog exhibits different modes of expression during embryogenesis. Development. 1993;119:673–90. http://www.ncbi.nlm.nih.gov/pubmed/8187637. Accessed 16 Mar 2018.

48. Li X, Zhao X, Fang Y, Jiang X, Duong T, Fan C, et al. Generation of destabilized green fluorescent protein as a transcription reporter. J Biol Chem. 1998;273:34970–5. http://www.ncbi.nlm.nih.gov/pubmed/9857028. Accessed 16 Mar 2018.

49. Novitch BG, Spicer DB, Kim PS, Cheung WL, Lassar AB. pRb is required for MEF2-dependent gene expression as well as cell-cycle arrest during skeletal muscle differentiation. Curr Biol. 1999;9:449–59. http://www.ncbi.nlm.nih.gov/pubmed/10322110. Accessed 16 Mar 2018.

50. Matus DQ, Lohmer LL, Kelley LC, Schindler AJ, Kohrman AQ, Barkoulas M, et al. Invasive Cell Fate Requires G1 Cell-Cycle Arrest and Histone Deacetylase-Mediated Changes in Gene Expression. Dev Cell. 2015;35:162–74. doi:10.1016/j.devcel.2015.10.002.

51. Nicolay BN, Bayarmagnai B, Moon NS, Benevolenskaya E V, Frolov M V. Combined inactivation of pRB and hippo pathways induces dedifferentiation in the Drosophila retina. PLoS Genet. 2010;6:e1000918. doi:10.1371/journal.pgen.1000918.

52. Xu Y, Swartz KL, Siu KT, Bhattacharyya M, Minella AC. Fbw7-dependent cyclin E regulation ensures terminal maturation of bone marrow erythroid cells by restraining oxidative metabolism. Oncogene. 2014;33:3161–71. doi:10.1038/onc.2013.289.

53. Mohamed TMA, Ang Y-S, Radzinsky E, Zhou P, Huang Y, Elfenbein A, et al. Regulation of Cell Cycle to Stimulate Adult Cardiomyocyte Proliferation and Cardiac Regeneration. Cell. 2018;173:104–116.e12. doi:10.1016/j.cell.2018.02.014.

54. Engerer P, Suzuki SC, Yoshimatsu T, Chapouton P, Obeng N, Odermatt B, et al. Uncoupling of neurogenesis and differentiation during retinal development. EMBO J. 2017;36:1134–46. doi:10.15252/embj.201694230.

55. Reis T, Edgar BA. Negative regulation of dE2F1 by cyclin-dependent kinases controls cell cycle timing. Cell. 2004;117:253–64. http://www.ncbi.nlm.nih.gov/pubmed/15084262. Accessed 28 Mar 2018.

56. Narasimha AM, Kaulich M, Shapiro GS, Choi YJ, Sicinski P, Dowdy SF. Cyclin D activates the Rb tumor suppressor by mono-phosphorylation. Elife. 2014;3. doi:10.7554/eLife.02872.

57. Ruijtenberg S, van den Heuvel S. G1/S Inhibitors and the SWI/SNF Complex Control Cell-Cycle Exit during Muscle Differentiation. Cell. 2015;162:300–13. doi:10.1016/j.cell.2015.06.013.

58. Albini S, Coutinho Toto P, Dall’Agnese A, Malecova B, Cenciarelli C, Felsani A, et al. Brahma is required for cell cycle arrest and late muscle gene expression during skeletal myogenesis. EMBO Rep. 2015;16:1037–50. doi:10.15252/embr.201540159.

59. Nègre N, Brown CD, Ma L, Bristow CA, Miller SW, Wagner U, et al. A cis-regulatory map of the Drosophila genome. Nature. 2011;471:527–31. doi:10.1038/nature09990.

60. Agawa Y, Sarhan M, Kageyama Y, Akagi K, Takai M, Hashiyama K, et al. Drosophila Blimp-1 is a transient transcriptional repressor that controls timing of the ecdysone-induced developmental pathway. Mol Cell Biol. 2007;27:8739–47. doi:10.1128/MCB.01304-07.

61. Akagi K, Sarhan M, Sultan A-RS, Nishida H, Koie A, Nakayama T, et al. A biological timer in the fat body comprising Blimp-1, βFtz-f1 and Shade regulates pupation timing in Drosophila melanogaster. doi:10.1242/dev.133595.

62. O’Keefe DD, Prober DA, Moyle PS, Rickoll WL, Edgar BA. Egfr/Ras signaling regulates DE-cadherin/Shotgun localization to control vein morphogenesis in the Drosophila wing. Dev Biol. 2007;311:25–39. doi:10.1016/j.ydbio.2007.08.003.

63. Etournay R, Popović M, Merkel M, Nandi A, Blasse C, Aigouy B, et al. Interplay of cell dynamics and epithelial tension during morphogenesis of the Drosophila pupal wing. Elife. 2015;4:e07090. doi:10.7554/eLife.07090.

64. Ma Y, Kanakousaki K, Buttitta L. How the cell cycle impacts chromatin architecture and influences cell fate. Front Genet. 2015;6. doi:10.3389/fgene.2015.00019.

65. Schulz KN, Bondra ER, Moshe A, Villalta JE, Lieb JD, Kaplan T, et al. Zelda is differentially required for chromatin accessibility, transcription factor binding, and gene expression in the early Drosophila embryo. Genome Res. 2015;25:1715–26. doi:10.1101/gr.192682.115.

66. Pajoro A, Madrigal P, Muiño JM, Matus JT, Jin J, Mecchia MA, et al. Dynamics of chromatin accessibility and gene regulation by MADS-domain transcription factors in flower development. Genome Biol. 2014;15:R41. doi:10.1186/gb-2014-15-3-r41.

67. de la Torre-Ubieta L, Stein JL, Won H, Opland CK, Liang D, Lu D, et al. The Dynamic Landscape of Open Chromatin during Human Cortical Neurogenesis. Cell. 2018;172:289–304. e18. doi:10.1016/j.cell.2017.12.014.

68. McKay DJ, Lieb JD. A common set of DNA regulatory elements shapes Drosophila appendages. Dev Cell. 2013;27:306–18. doi:10.1016/j.devcel.2013.10.009.

69. Langmead B, Salzberg SL. Fast gapped-read alignment with Bowtie 2. Nat Methods. 2012;9:357–9. doi:10.1038/nmeth.1923.

70. Zhang Y, Liu T, Meyer CA, Eeckhoute J, Johnson DS, Bernstein BE, et al. Model-based analysis of ChIP-Seq (MACS). Genome Biol. 2008;9:R137. doi:10.1186/gb-2008-9-9-r137.

71. Zhang Y, Lin Y-H, Johnson TD, Rozek LS, Sartor MA. PePr: a peak-calling prioritization pipeline to identify consistent or differential peaks from replicated ChIP-Seq data. Bioinformatics. 2014;30:2568–75. doi:10.1093/bioinformatics/btu372.

72. Ramírez F, Ryan DP, Grüning B, Bhardwaj V, Kilpert F, Richter AS, et al. deepTools2: a next generation web server for deep-sequencing data analysis. Nucleic Acids Res. 2016;44:W160–5. doi:10.1093/nar/gkw257.

73. Robinson JT, Thorvaldsdóttir H, Winckler W, Guttman M, Lander ES, Getz G, et al. Integrative genomics viewer. Nat Biotechnol. 2011;29:24–6. doi:10.1038/nbt.1754.

74. Zhu LJ, Christensen RG, Kazemian M, Hull CJ, Enuameh MS, Basciotta MD, et al. FlyFactorSurvey: a database of Drosophila transcription factor binding specificities determined using the bacterial one-hybrid system. Nucleic Acids Res. 2011;39:D111–D117. doi:10.1093/nar/gkq858.

75. Bailey TL, Boden M, Buske FA, Frith M, Grant CE, Clementi L, et al. MEME SUITE: tools for motif discovery and searching. Nucleic Acids Res. 2009;37 Web Server issue:W202–8. doi:10.1093/nar/gkp335.

76. Zhu LJ, Gazin C, Lawson ND, Pagès H, Lin SM, Lapointe DS, et al. ChIPpeakAnno: a Bioconductor package to annotate ChIP-seq and ChIP-chip data. BMC Bioinformatics. 2010;11:237. doi:10.1186/1471-2105-11-237.

77. Dobin A, Davis CA, Schlesinger F, Drenkow J, Zaleski C, Jha S, et al. STAR: ultrafast universal RNA-seq aligner. Bioinformatics. 2013;29:15–21. doi:10.1093/bioinformatics/bts635.

78. Anders S, Pyl PT, Huber W. HTSeq--a Python framework to work with high-throughput sequencing data. Bioinformatics. 2015;31:166–9. doi:10.1093/bioinformatics/btu638.

79. Trapnell C, Williams BA, Pertea G, Mortazavi A, Kwan G, van Baren MJ, et al. Transcript assembly and quantification by RNA-Seq reveals unannotated transcripts and isoform switching during cell differentiation. Nat Biotechnol. 2010;28:511–5. doi:10.1038/nbt.1621.

80. Huang DW, Sherman BT, Lempicki RA. Systematic and integrative analysis of large gene lists using DAVID bioinformatics resources. Nat Protoc. 2009;4:44–57. doi:10.1038/nprot.2008.211.

81. Love MI, Huber W, Anders S. Moderated estimation of fold change and dispersion for RNA-seq data with DESeq2. Genome Biol. 2014;15:550. doi:10.1186/s13059-014-0550-8.

## Supplemental References

Djiane, A., Krejci, A., Bernard, F., Fexova, S., Millen, K., and Bray, S.J. (2013). Dissecting the mechanisms of Notch induced hyperplasia. EMBO J 32, 60–71.

Korenjak, M., Anderssen, E., Ramaswamy, S., Whetstine, J.R., and Dyson, N.J. (2012). RBF binding to both canonical E2F targets and noncanonical targets depends on functional dE2F/dDP complexes. Mol Cell Biol 32, 4375–4387.

Oh, H., Slattery, M., Ma, L., Crofts, A., White, K.P., Mann, R.S., and Irvine, K.D. (2013). Genome-wide association of Yorkie with chromatin and chromatin-remodeling complexes. Cell Rep 3, 309–318.

